# Co-activation of LIN28A and CTNNB1 disturbs cortical neuronal migration and pia mater integrity

**DOI:** 10.1101/2024.08.01.606182

**Authors:** Jelena Navolić, Sara Hawass, Manuela Moritz, Jan Hahn, Maximilian Middelkamp, Antonia Gocke, Matthias Dottermusch, Yannis Schumann, Lisa Ruck, Christoph Krisp, Shweta Godbole, Piotr Sumislawski, Hartmut Schlüter, Julia E. Neumann

## Abstract

Developmental signalling pathways act in stage and tissue dependent relation and mis-activation can drive tumour formation. The RNA-binding protein LIN28A maintains stemness and is overexpressed in embryonal brain tumours. Activating mutations of *CTNNB1* - the WNT pathway effector - have been reported in respective brain tumours. The aim of this study was to investigate the interplay of these oncogenic proteins during embryonal brain development.

The combination of both oncogenic factors did not lead to brain tumour formation but resulted in disturbed lamination and impaired cell migration in the cerebral cortex. Spatially resolved proteome analysis revealed imbalances of the extracellular matrix protein LAMB1 and its receptors RPSA and ITGB1 accompanied by a porous pial border and overmigration of neural cells. Cajal-Retzius cells were misplaced in deeper cortex regions without affecting general REELIN levels and additional reduced levels of α-DYSTROGLYCAN.

Taken together, the interplay of LIN28A and CTNNB1 resulted in a cortical migration disorder showing histomorphological and molecular similarities to human Cobblestone lissencephaly (type 2), highlighting novel implications of the oncogene LIN28A in extracellular matrix integrity.

## Introduction

Cortical brain development is a process that requires precisely controlled spatial and temporal signalling. One such process is the development of the six-layered cerebral cortex (Cooper, 2008) including the generation of neural cells (Götz and Huttner, 2005), differentiation (Noctor *et al*., 2004) and radial migration (Rakic, 1972; Nadarajah and Parnavelas, 2002). Disturbances of this processes can lead to a variety of developmental disorders (Hevner, 2007) and to tumorigenesis (Ben-Porath *et al*., 2008; Pajtler *et al*., 2015; Jessa *et al*., 2019; Hovestadt *et al*., 2020; Curry and Glasgow, 2021). One pathway, which plays a key role in processes of brain development, is the WNT-pathway (Harrison-Uy and Pleasure, 2012). In canonical WNT-pathway activation, the WNT ligand binds to its receptor FRIZZLED and the downstream effector protein CTNNB1 enters the nucleus and acts as a transcriptional factor (Komiya and Habas, 2008). WNT-pathway activation is described in early childhood brain tumours, such as in highly aggressive embryonal tumours with multi-layered rosettes (ETMRs) arising predominantly in the forebrain. One of the ETMR hallmarks - also used in diagnostics - is the overexpression of LIN28A (Korshunov *et al*., 2012; Lambo *et al*., 2019, 2020). LIN28A is an RNA-binding protein and a known inhibitor of the let-7 miRNA family, whose components are acting as tumour suppressors (Li and He, 2012; Thornton and Gregory, 2012). By inhibition of the mi-RNA family, LIN28A maintains proliferation and self-renewal of stem cells during early development (Viswanathan *et al*., 2009; Zhong *et al*., 2010; Balzeau *et al*., 2017; Herrlinger *et al*., 2019). The mechanisms leading to an upregulation of LIN28A during tumorigenesis and its specific oncogenic function is not yet fully understood (Viswanathan *et al*., 2009). The lacking comprehensive knowledge about ETMR tumour biology impedes the finding appropriate therapy targets and affected children face a poor prognosis (Pollack, Agnihotri and Broniscer, 2019). It was shown that GFAP^+^ neural precursor cells in the ventricular zone (VZ) of the cerebral cortex are potential cells of origin for ETMR (Neumann *et al*., 2017; Lambo *et al*., 2019, 2020). These neural precursor cells expand first their pool of cells, and then differentiate into radial glial cells (RGs) establishing a scaffold for radial migration of neuronal cells to their anticipated locations in the cerebral cortex (Rakic, 1972; Nadarajah and Parnavelas, 2002). An *in vivo* model with the sole overexpression of LIN28A in GFAP^+^ neural precursor cells does not lead to tumour formation but results in a transiently higher proliferation rate during embryonic development (Middelkamp *et al*., 2021). The sole activation of the WNT-pathway in GFAP^+^ neural precursor cells was neither sufficient to drive tumorigenesis but led to a disturbance in cortical lamination (Pöschl *et al*., 2013). The interplay of the WNT pathway with another known developmental pathway – the sonic hedgehog (SHH)-pathway – resulted in the formation of ETMR-like tumours. However, these do not show LIN28A expression, implying that the upregulation of LIN28A precedes the WNT- and SHH-pathway activation to facilitate tumour growth (Neumann *et al*., 2017).

The aim of this study was to investigate if co-activation of LIN28A and the WNT pathway in neural precursor cells affect brain development and are sufficient and necessary to initiate tumour formation. We, therefore, generated mouse models expressing each factor alone, or both in combination in hGFAP^+^ precursors and analysed brain morphology and spatial molecular profiles. As LIN28A and the WNT pathway have important regulatory functions in transcription and translation, our focus was to investigate the protein composition of the cerebral cortex with spatial resolution and to examine structural and functional changes in the forebrain region. We show that the interplay of LIN28A and CTNNB1 was not sufficient to initiate tumour growth during embryonal brain development but result in spatial disturbances of components of the extracellular matrix and morphological changes associated with a lissencephaly type 2 -like phenotype.

## Results

### Disturbance of cortical lamination but no tumour formation

To analyse the interplay of LIN28A and WNT signalling during embryonal brain development, we generated mice displaying either sole overexpression of LIN28A (GL), a sole overexpression of stabilised CTNNB1 (GB) or the combination of both (GBL) in hGFAP^+^ neural precursor cells (Appendix Fig. S1). We evaluated brain histomorphology at embryonic (E) time points E14.5 (shortly after *hGFAP- cre* promotor activation (Zhuo *et al*., 2001)) and E18.5 (shortly before birth, (Fig. 1). Co-activation of both factors did not lead to brain tumour formation (Fig. 1). In line with previous results (Middelkamp *et al*., 2021), the GL model did not show coarse morphological changes compared to the CTRL. In contrast, GB and GBL models developed a hydrocephalus and strong disturbances of cortical lamination already apparent at E14.5. At E18.5 isocortical layering could not be detected in GB and GBL mice. The GBL model additionally displayed variable cortex thickness and large blood vessels in deeper regions around E18.5 (indicated by arrows, Fig. 1) with lethal consequences around the birth time. The cortical marker SOX2 is expressed in the ventricular zone (VZ) during development (Miyagi *et al*., 2006; Amador-Arjona *et al*., 2015; Penisson *et al*., 2019) (Appendix Fig. S2). In GBL mice this primary proliferative zone was generally established but highly disturbed compared to the other conditions (Appendix Fig. S2). In both, the GB and GBL models, we observed a lack of TBR2^+^ intermediate progenitor cells (IPs) of the subventricular zone (SVZ) (Wrobel *et al*., 2007). Expression of the somatostatin receptor SSTR2 (Appendix Fig. S2) in migrating cells leaving the ventricular zone (Le Verche *et al*., 2009; Bedogni and Hevner, 2021) was disturbed in the SVZ and intermediate zone (IZ) with patchy appearance in the GBL model. NEUN, a marker for differentiated neuronal cells (Gusel’nikova and Korzhevskiy, 2015) showed positive cells in the cortical plate at E18.5 in the CTRL and GL model. The GB and GBL model displayed positive cells but in random distribution with no clear areal specification. These results demonstrate that LIN28A and CTNNB1 are not sufficient to drive early tumour formation in the brain but result in disturbed formation of neural cells and disordered lamination.

**Fig 1:**
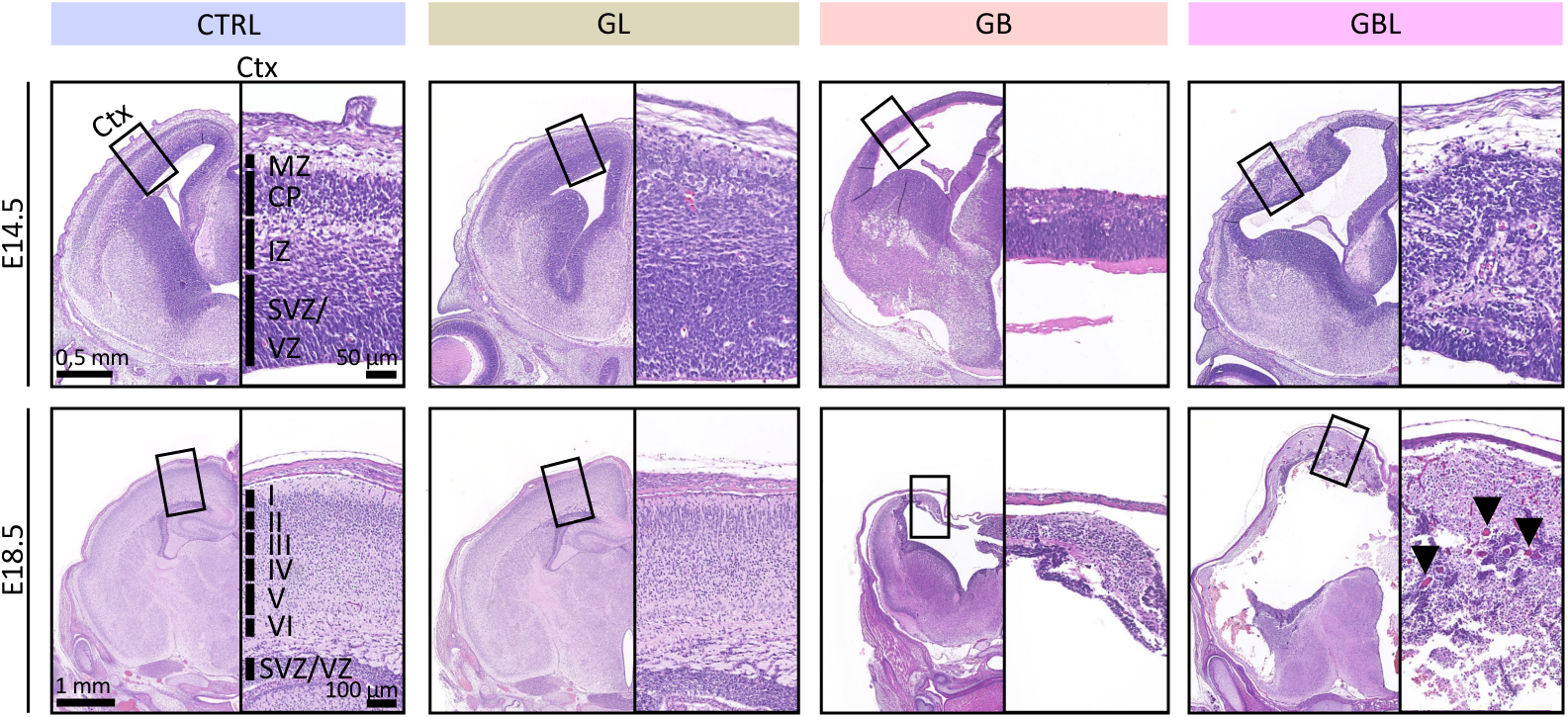
**Co-expression of LIN28A and stabilised CTNNB1 results in severe changes of forebrain histomorphology**. H&E-stained frontal forebrain sections of control mice (CTRL), transgenic mice overexpressing LIN28A (GL), mice overexpressing stabilised CTNNB1 (GB) and mice co-expressing both factors (GBL) at embryonal days E14.5 and E18.5. The embryonic head (left) and a magnification of the cerebral cortex (Ctx, right) is shown for each mouse model. GB mice display a hydrocephalus and thinned cerebral cortex. GBL mice show severely disturbed cortical lamination and a variable cortex thickness. Arrows indicate larger blood vessels in the brain parenchyma of GBL mice, which are not present in the other models. MZ = marginal zone, CP = cortical plate, IZ = intermediate zone, SVZ = subventricular zone, VZ = ventricular zone, I-VI = cortex layer I-VI.

### Disturbed cortical migration and proliferation

We hypothesised that disturbed cortical lamination might result from impaired neural migration during cortical brain development. We, therefore, performed cell tracking of BrdU^+^ cells between day E14.5 and E16.5 during embryonic development (Fig. 2A). The mean cerebral cortex thickness was significantly thinner in the GB and GBL models within the respective regions of interest (CTRL: x̅ = 456.83 µm, GL: x̅ = 447.38 µm, GB: x̅ = 240.92 µm, GBL: x̅ = 225.12 µm, n = 3, p < 0.05, Fig. 2C) and significantly less BrdU^+^ cells in total were found (CTRL: x̅ = 607, GL: x̅ = 545, GB: x̅ = 380, GBL: x̅ = 371, p < 0.05). Distance quantification from the ventricular margin (0 µm) to each BrdU^+^ cell in the cerebral cortex (Fig. 2B) revealed a significantly reduced mean migration distance in the GB and GBL models compared to the control (CTRL) and GL model (CTRL: x̅ = 171.94 µm, GL: x̅ = 182.90 µm, GB: x̅ = 107.15 µm, GBL: x̅ = 112.85 µm, n = 3, p < 0.05, Fig. 2C). In the GB and GBL models, a significantly smaller number of BrdU^+^ cells are located more distant than 200 µm and no cells were observed to migrate further than 400 µm within the region of interest (Fig. 2D).

**Fig 2:**
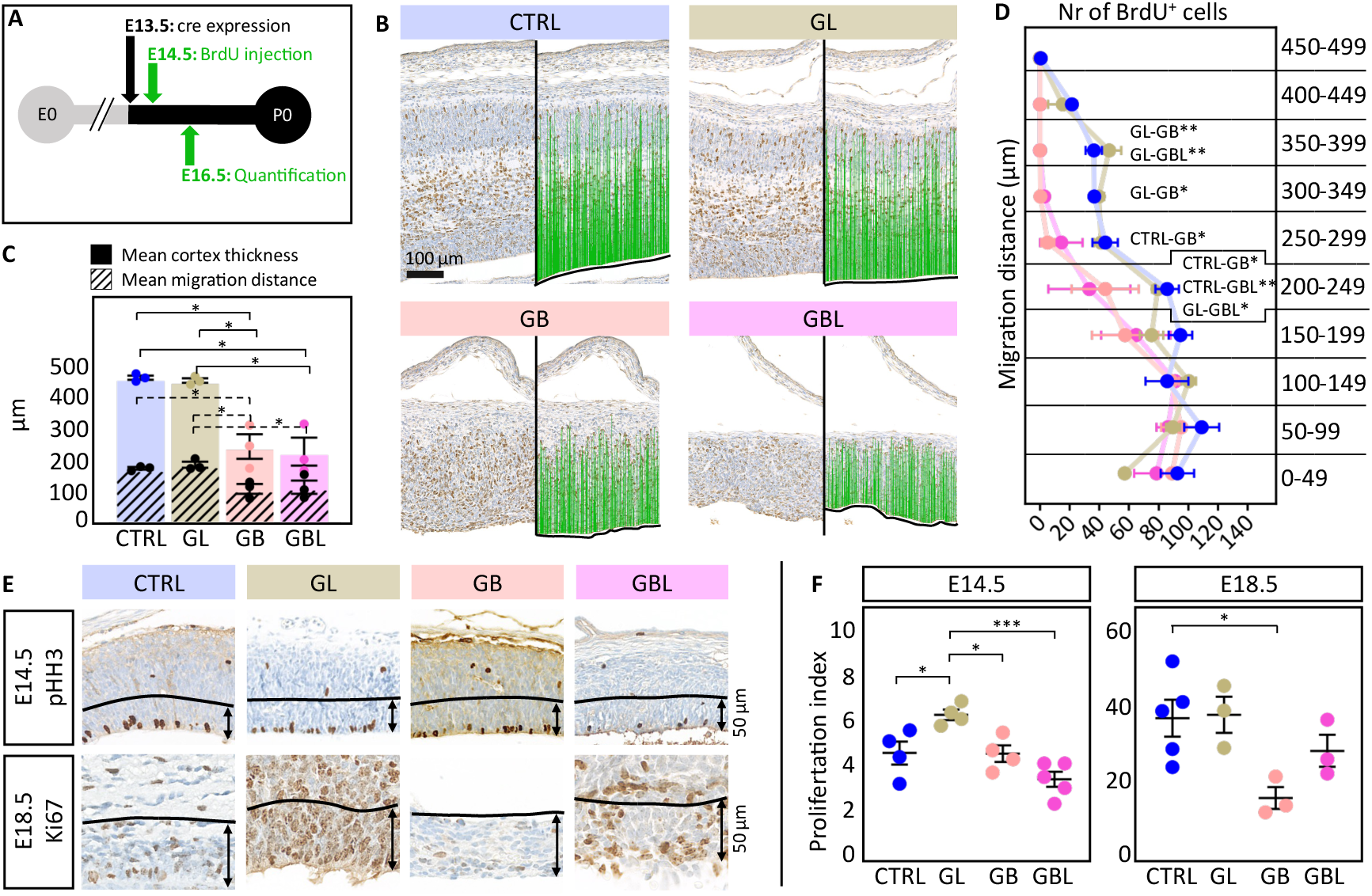
Alterations of cortical cell migration and proliferation during cerebral development in GL, GB and GBL mice. **(A)** Experimental design of BrdU (Bromodeoxyuridine) injection and quantification time points. **(B)** BrdU staining of frontal sections of cerebral cortices at E16.5 (left) and the respective distance quantification output (right): blue lines indicate the migration distance of BrdU^+^ cells from the ventricular margin (dashed line). **(C)** Mean cortex thickness and mean migration distance across mouse models. Significant differences between models are indicated. Colour filled bars indicate mean cortex thickness and dashed bars indicate mean migration distance (n = 3, one-way ANOVA). **(D)** Number of BrdU^+^ cells per 50 µm distance bins ranging from 0 to 500 µm across the models. Significant migration differences between models for the respective bins are indicated (n = 3, two-way ANOVA). **(E)** Immunohistochemical staining of frontal sections of cerebral cortices against proliferation markers pHH3 (E14.5, upper panels) and ki67 (E18.5, lower panels). Arrows indicate the area (spanning 50 µm from the ventricular border) used for quantification of positively stained cells in **(F). (F)** Proliferation index indicates the number of pHH3 or Ki67^+^ cells per 100 µm (n >= 3, one-way ANOVA). For C, D, F: * p < 0.05, ** p < 0.01, *** p < 0.001, mean and SEM are shown.

Previous studies have shown increased transient proliferation in the GL model (Middelkamp *et al*., 2021) and a decrease in the GB model (Pöschl *et al*., 2013) during embryonic development. Here, the proliferation rate (cells^+^/100µm, Fig. 2F) was determined based on immunohistochemically stained cortex sections using antibodies against phosphorylated histone H3 (pHH3) (E14.5) and Ki67 (E18.5) (Fig. 2E). As expected, a transiently higher proliferation rate at E14.5 was observed in the GL model which was abolished at E18.5, and a reduced proliferation rate in the GB model at E18.5. For the GBL model - despite a histologically disturbed ventricular zone (Fig. 2E, Appendix Fig S2) - no significant difference in proliferation was detected (Fig. 2F). In conclusion, activation of CTNNB1 resulted in impaired neural migration and decreased proliferation during development. Co-activation of LIN28A re-established proliferation rates within the cortex, however, a severe migration deficit was still apparent.

### Spatially resolved proteomics of the developing brain

In order to elucidate the molecular mechanisms resulting in the observed phenotypes we aimed to analyse molecular parameters in a spatially resolved manner. As LIN28A and CTNNB1 are both transcriptional and translational regulators, we aimed to investigate the resulting protein level using mass spectrometry-based proteome data with high spatial resolution. Therefore, we used the nanosecond infrared laser (NIRL) nano-volume sampling system (Hahn *et al*., 2021; Voß *et al*., 2022) and ablated consecutive cortical layers directly from the skin surface into the cerebral cortex, as described previously (Navolić *et al*., 2023). We ablated nine consecutive layers at E14.5 and 18 consecutive layers at E18.5 of the CTRL condition in biological replicates (n = 4) and the mouse models GL, GB and GBL (n = 3) (Fig. 3A, Appendix Fig. S3A). The layers were ablated in four batches (LA.batch) resulting in a data set with in total 367 samples measured in eight batches (m.batch) and identifying over 5000 proteins (Fig. 3B). The initial sample distribution was mainly driven by developmental differences according to the time points (Appendix Fig. S3B). Hence, downstream analysis was done separately for E14.5 (124 samples) and E18.5 (243 samples) (Appendix Fig. S3C). By applying the batch effect reduction algorithm BERT ((Schumann and Schlumbohm, 2024) in preparation), we reduced technical batch effects and preserved over 3000 proteins providing quantitative information for comprehensive analysis. Dimension reduction analysis based on 100% valid values showed a gradient distribution from superficial to deeper layers by the represented proteins (Fig. 3C). To better understand similarity across layers, we performed consensus clustering based on the CTRL samples and detected two clear layer clusters at E14.5 (C_1_ and C_2_) and three clusters at E18.5 (C_3_ - C_5_) (Fig. 3D, Appendix Fig. S4A). Marker proteins for the skin (Uhlén *et al*., 2015), bone (Janssen *et al*., 2023), meningeal structures (Uhlén *et al*., 2015; DeSisto *et al*., 2020) and cerebral cortex (Uhlén *et al*., 2015) allowed to determine anatomical identities of the ablated layers (Fig. 3E). Layers belonging to the clusters C_1_, C_3_ and C_4_ showed higher abundances of proteins associated with the skin, bone, and meningeal structures (Fig. 3E). The clusters C_2_ and C_5_ showed higher abundances of cerebral cortex markers (Fig. 3E) and were differently expressed across the GL, GB and GBL layers (Appendix Fig. S4B). Taken together spatially resolved proteomics distinguished cortical layers and revealed disturbances in the GL, GB and GBL models, as compared to the CTRL.

**Fig 3:**
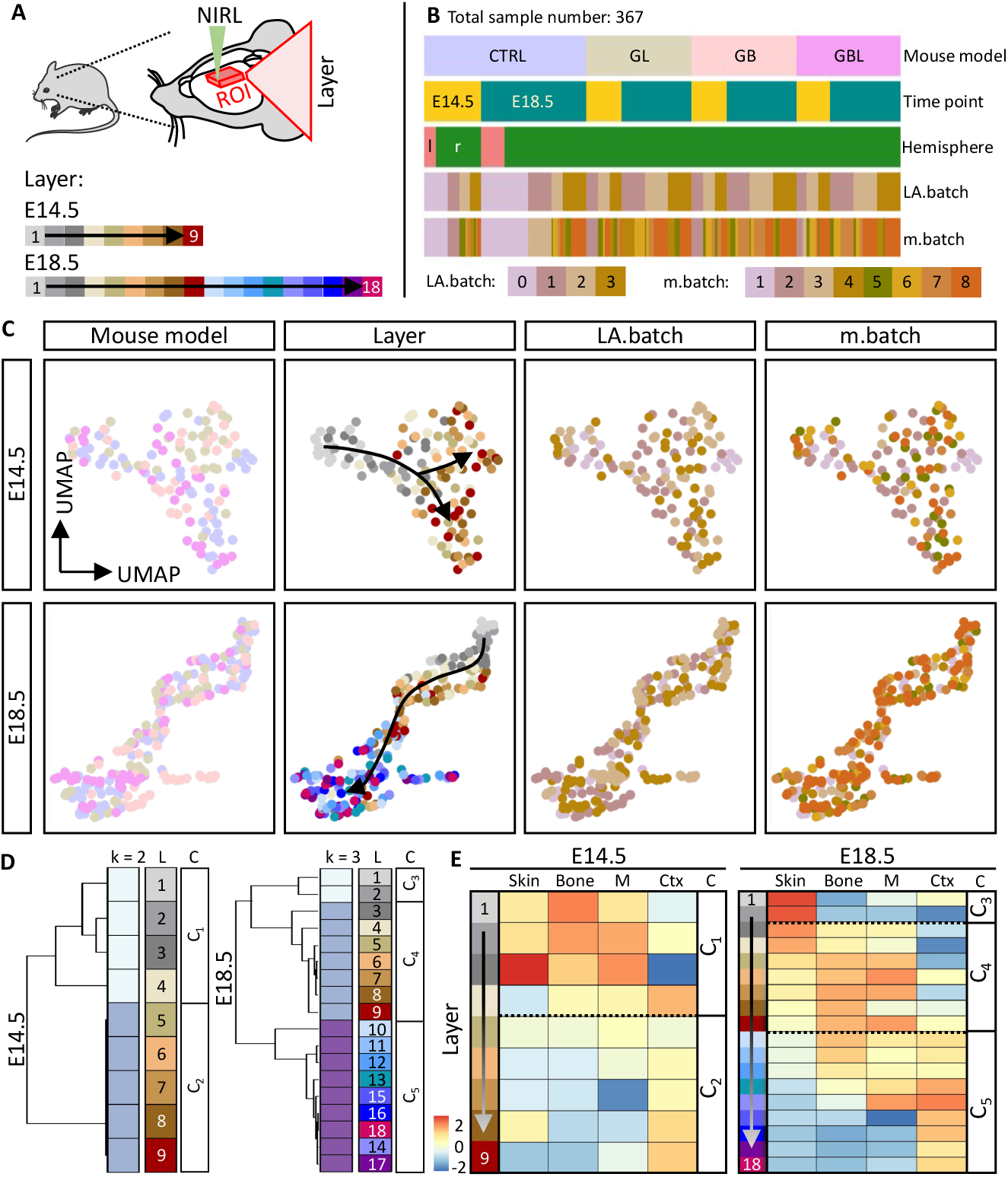
Spatial proteome analysis of the murine cortex of CTRL, GL, GB and GBL mice. **(A)** Experimental design for spatial proteomics using the nanosecond infrared laser (NIRL) ablation system to ablate consecutive layers (∼40 µm) directly from the frozen sample targeting the region of interest (ROI) within the forebrain region as previously described (Navolić et al. 2023). At E14.5 nine consecutive layers were ablated and at E18.5 18 layers were ablated. **(B)** In total 367 samples were obtained, including CTRL (n = 4), GL (n = 3), GB (n = 3) and GBL (n = 3) for each time point E14.5 (124 samples) and E18.5 (243 samples). The samples were ablated and further processed in four batches (LA.batch) and mass spectrometric measurements were performed in eight batches (m.batch). **(C)** Uniform Manifold Approximation and Projection (UMAP) for the E14.5 and E18.5 dataset based on proteins with 100% valid values. Legends in **(A)** and **(B)** apply for colour scheme in **(C)**. **(D)** Consensus clustering analysis combines individual layers into layer clusters: C_1_ and C_2_ for E14.5 and C_3_ - C_5_ for E18.5. k = number of clusters, L = layer, C = cluster. **(E)** Mean abundance (column scaled) of marker proteins for skin (FLG, KRT14, LORICRIN; (Uhlén et al. 2015)), bone (COL1A1, COL1A2, SERPINF1; (Janssen et al. 2023)), meninges ( = M; CDH11, CRABP2, TAGLN; (DeSisto et al. 2020; Uhlén et al. 2015)) and cerebral cortex ( = Ctx; TBR1, MAP2, BCL11B; (Uhlén et al. 2015)). C = cluster C_1_ - C_5_ are indicated with dashed lines.

### Characterisation of unique layer signatures

We further characterised a unique signature for each CTRL layer based on the unique top 10 high abundant proteins for each layer (Table 4). Correlation analysis among the CTRL samples demonstrates that these unique top 10 high abundant proteins are sufficient for layer characterisation (Fig. 4A, B, D, E, Appendix Fig. S5A). The top 10 high abundant proteins were detected in all mouse models, but the characteristic pattern observed in the CTRL condition was abolished among the GL, GB and GBL layers (Appendix Fig. S5B, C, D, E). To determine the highest similarity of cortical layers in the GL, GB and GBL models compared to the CTRL, we selected the highest correlation value of each layer comparison (Appendix Fig. S5C, E). Compared to CTRL, most layers of GL mice fell into similar layers or the same layer cluster (C_1_-C_5_, Appendix Fig. S5B-E). In contrast, deeper layers in the GB model (E14.5, x-axis: C_2_) were more similar to superficial CTRL layers (y-axis: C_1_). In the GBL model (E18.5) the superficial layers (x-axis: C_1_ and C_2_) resembled deeper CTRL layers belonging to C_4_ and C_5_ (y-axis) (Fig. 4C, F).

**Fig 4:**
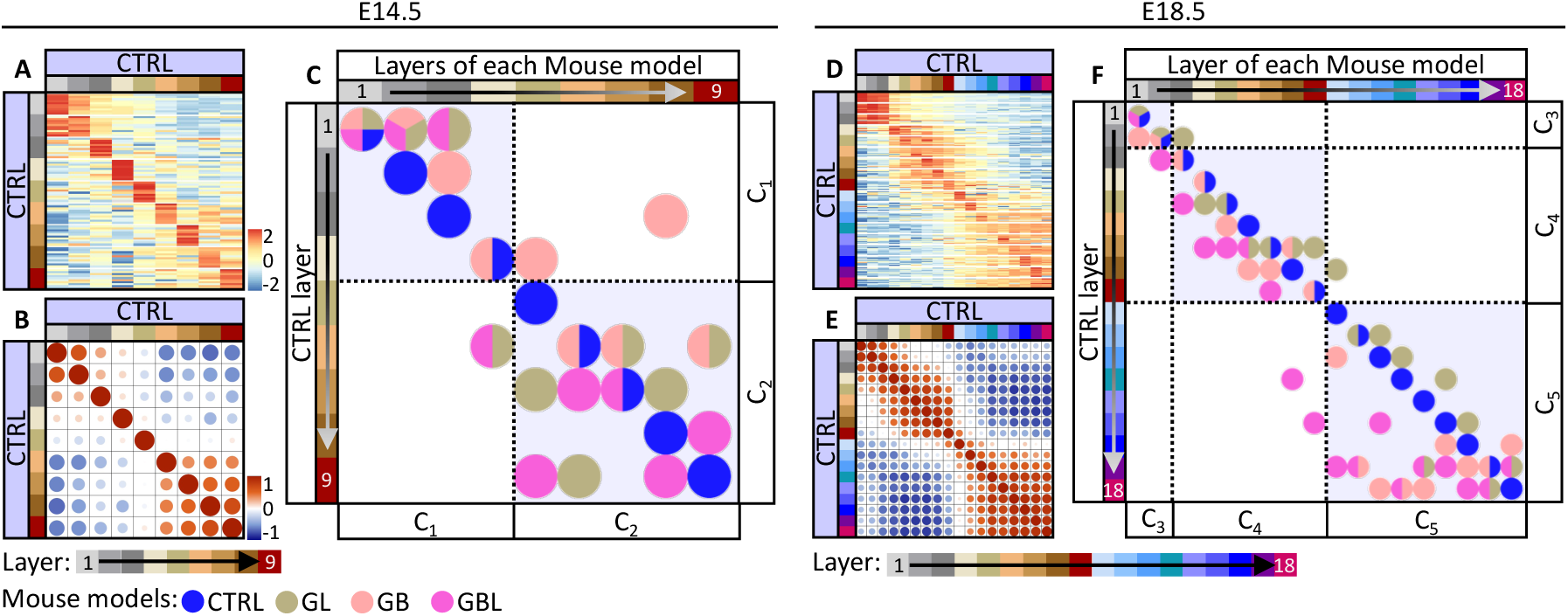
Spatial representation of CTRL layer signature proteins across mouse models. **(A, D)** Heatmap representing the top 10 high abundant proteins (Table 4) of each CTRL layer (y-axis) in all CTRL layers (x-axis) at E14.5 **(A)** or E18.5 **(D)**. Scaled rows**. (B, E)** Correlation analysis based on the top 10 high abundant proteins of each CTRL layer at E14.5 **(B)** or E18.5 **(E)**. **(C, F)** Summary of correlation analysis results based on the top 10 high abundant proteins of each CTRL layer. Dots represent the highest positive correlation for each CTRL, GL, GB and GBL layer (x-axis) compared to the CTRL layer (y-axis). Layer clusters C_1_-C_5_ are indicated with dashed lines. At E18.5 superficial GBL layers show a shift of protein patterns towards deeper layers of the CTRL.

Generally, all mouse models had a characteristic unique layer signature capturing the differences across layers (Appendix Fig. S6, Table 4). Gene ontology analysis for biological processes (condensed into parent GO:BP-terms to reduce redundant terms, Table 5) indicated functions that are represented similarly across the models (E18.5: “skin development”). Other functions appear rather in the recombinant models GL, GB and GBL (E14.5: “gene expression”). Some terms show a greater shift of the representing layers from superficial layers in the CTRL to the deeper layers in the GBL model (E14.5: “epidermis development”; E18.5: “cellular response to monoamine stimulus”, Appendix Fig. S6). In summary, the GB and GBL models showed differences in spatial protein abundances. The GBL displayed a shift in layer identity with superficial layers being more similar to deeper layers of the CTRL.

### Imbalanced protein abundances associated with a lissencephaly-like phenotype

In order to capture specific changes in protein abundance in a spatial context, we performed analysis of covariance (ANCOVA) for each layer cluster separately (C_1_ – C_5_) and compared the CTRL conditions with the other mouse models (GL, GB or GBL, Fig. 5A). Specific proteins for GL, GB or GBL with a significantly different rate of change (slope) or abundance (intercept) compared to the CTRL (p < 0.05 and R^2^ > 0.1, Fig. 5A, B) in either cluster were further described using gene ontology analysis for biological processes (GO:BP-terms, Fig. 5C). In the GL and GBL model, terms of transcription and translation appeared but were associated with different proteins. The GB model was defined by distinct GO:BP-terms, such as cell polarity and cellular component assembly (Fig. 5C). Of note, the GL and GBL models had a high proportion of altered proteins that were reported as direct interaction partners of LIN28A at protein or mRNA level, potentially affecting their translation (>40%, Fig. 5C) (Hafner *et al*., 2013; Parisi *et al*., 2021). We further categorised the proteins using a literature mining tool (modified version (Steffen *et al*., 2020)) to highlight potential proteins of interest (green colour code in Fig. 5C distinguishing between categories: 1 = at least one review paper/well known, 2 = at least one research paper/known, 3 = no further literature available/not known in context of “cortical development” as keyword (Steffen *et al*., 2020)). Beyond others, the laminin receptor protein RPSA was significantly altered in the GBL model. RPSA was shown to be involved in radial glia morphology shaping and affects cortical migration (Blazejewski *et al*., 2022). We therefore analysed the spatial resolved abundances of RPSA in more detail and further focused on known associated factors like Laminin (LAMB1), the ligand Serpinf1 (PEDF) (Blazejewski *et al*., 2022) and Integrins (ITGB1, Fig. 5D). The factors showed a strong deviation in the GBL model. We particularly observed a striking drop of LAMB1 in the C_1_ and C_4_ layers - representing meningeal structures - within the GBL model, but not in the other models (Fig. 5D).

**Fig 5:**
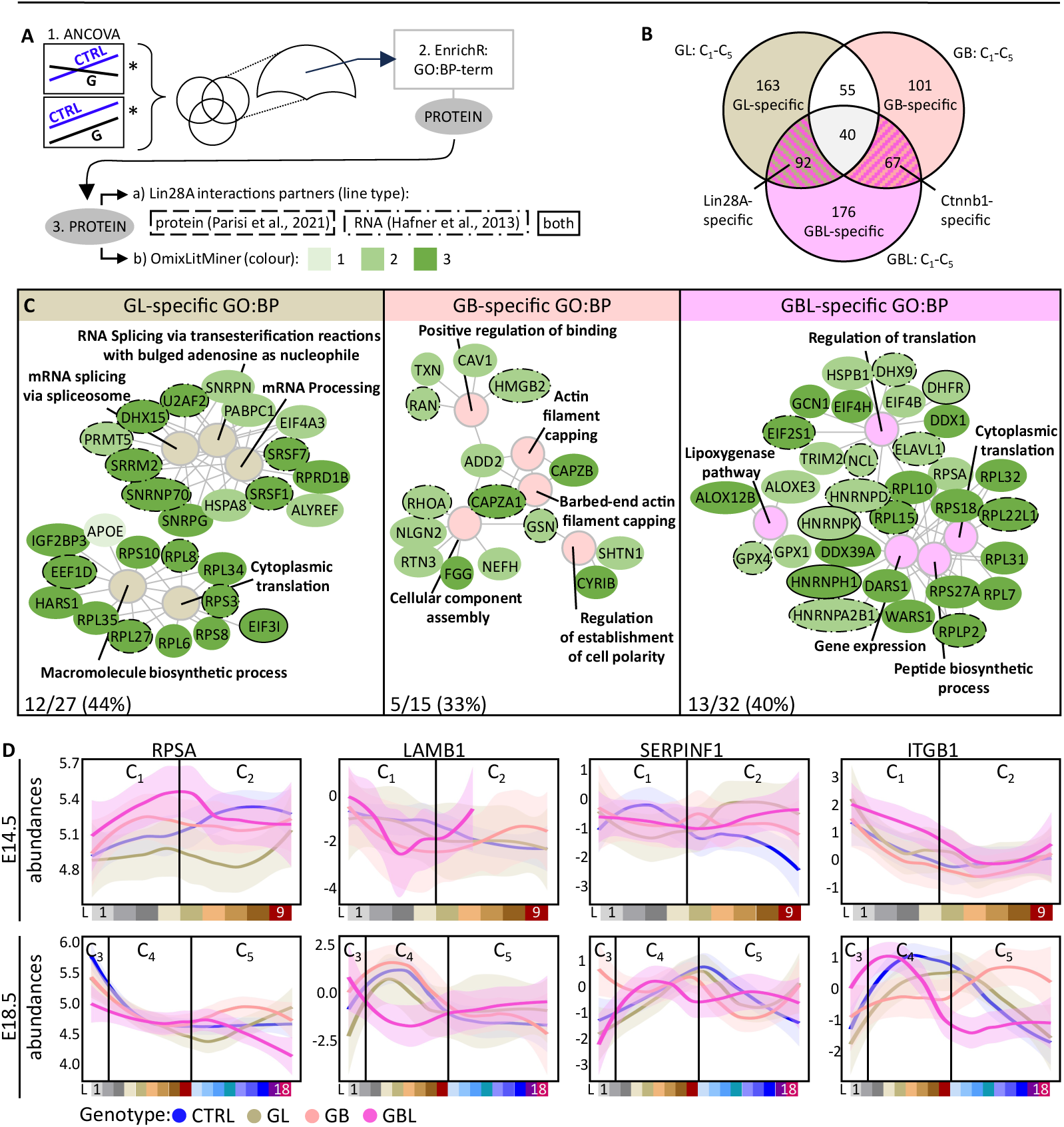
Layer and mouse model specific protein representation reveals ribosomal proteins and extracellular matrix components to be differentially distributed in brains with combined CTNNB1 and LIN28A activation. **(A)** Workflow analysing proteins of interest considering spatial resolution and mouse models. Proteins with significant difference upon ANCOVA based regression analyses (*) were taken as specific proteins for each mouse model after generating a Venn diagram. These proteins were further analysed using EnrichR (E. Y. Chen et al. 2013; Kuleshov et al. 2016; Xie et al. 2021) for gene ontology terms for biological processes (GO:BP-term). The associated proteins were then categorised by a) known LIN28A interaction patterns (indicated by line type) and b) the background knowledge using a literature mining tool (modified version (Steffen et al. 2020)) using the keyword “cortical development” and its respective categories: 1 = at least one review paper, 2 = at least one research paper, 3 = no further literature available. **(B)** Venn diagram of significant proteins after ANCOVA between CTRL vs GL or GB or GBL for all layer clusters (C_1_-C_5_). **(C)** Top 5 GO:BP-terms for each model-specific protein collection. Proteins are coloured based on the OmixLitMiner categories. The line types indicate previously described interaction with LIN28A. The percentage shows the proportion of potential partners to the total number of proteins for each panel (bottom corner). **(D)** Spatially resolved protein abundances of extracellular matrix components and their respective receptors. L = layer, C_1_-C_5_ = defined layer clusters based on consensus clustering analysis in CTRLs.

Immunohistochemical investigation of LAMB1/LAMA1 expression revealed a porous pial border in the GBL model at (E14.5) with aggregations of neuronal tissue (MAP2 positive) (Caceres *et al*., 1984) above that border (Fig. 6A). At E18.5, no clear pial border could be observed in GBL with larger blood vessels in deeper cortical regions (Fig. 1, Fig. 6A). The overmigration phenotype observed in the GBL model resembles the phenotype of the human migration disorder lissencephaly type 2, which is associated with hypo-glycosylation of α-DYSTROGLYCAN (α-DYS) (Barresi and Campbell 2006). We therefore analysed α-DYS levels and detected a significant reduction in the CTNNB1 associated GB and GBL models at E18.5, but no significant difference in the β-DYSTROGLYCAN (Fig. 6B, C, Appendix Fig. S7A). The impaired and disturbed cortical migration phenotype of the GB model is similar to the phenotype of the human disease lissencephaly type 1, which is characterised by reduction of REELIN, an important regulator of radial migration, mainly expressed by Cajal-Retzius cells (CRs) (Hirotsune *et al*., 1995; D’Arcangelo and Curran, 1998; Lambert de Rouvroit and Goffinet, 2001; Causeret *et al*., 2021). At E14.5 REELIN expression was significantly reduced in the GB model compared to the GL and GBL model with elevated REELIN abundance (p < 0.05, Fig. 6B, C). Histological observation showed ectopically located CRs in deeper layers of the cortex in the GBL model (Fig. 6C, Appendix Fig. S7C). In summary CTNNB1 activation resulted in a lissencephaly type 1-like phenotype with reduction of REELIN levels, whereas a co-activation of both LIN28A and CTNNB1 rescued the REELIN levels but led to a porous pial border resulting in a lissencephaly type 2- like phenotype.

**Fig 6:**
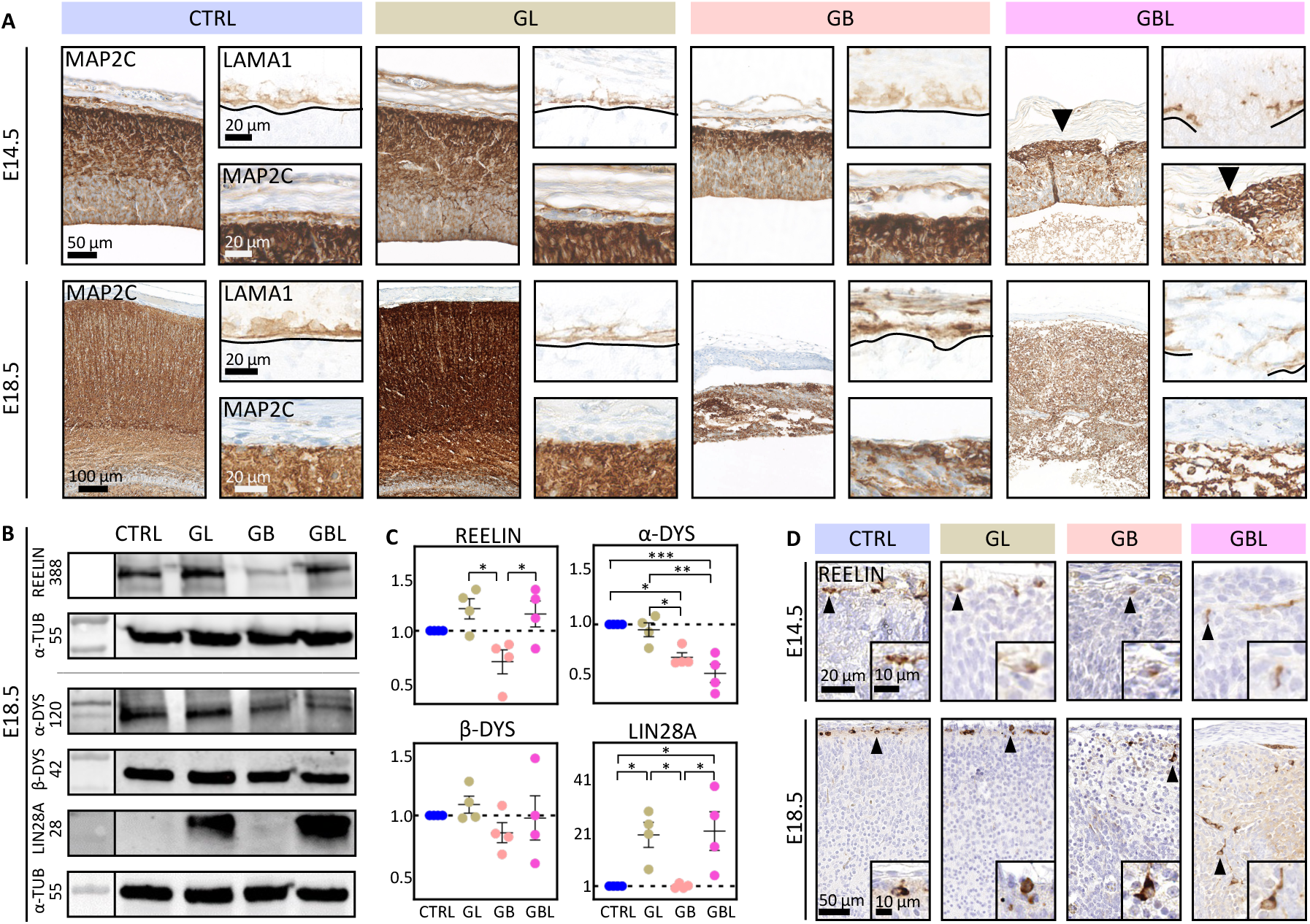
Porous pial border and neural overmigration in GBL mice. **(A)** Immunohistochemistry against LAMININ (LAMA1) as pial marker, and against MAP2C as dendritic marker for neuronal tissue. Pial border is indicated by dashed line. Ectopic neuronal tissue located above the pial border in the GBL model is indicated by arrows**. (B)** Western blot of cortical lysates stained against REELIN (388 kDa), α-DYS (α-DYSTROGLYCAN, 120 kDa), β-DYS (β-DYSTROGLYCAN, 42 kDa) and LIN28A (28 kDa) at E18.5. α-TUB (α-TUBULIN, 55 kDa) was used as housekeeping protein for normalisation. **(C)** Respective quantification of western blot signals **(B)** based on normalised values (n >=3, one-way ANOVA, * p < 0.05, ** p < 0.01, *** p < 0.001; mean and SEM are shown). REELIN levels were decreased in the GB model compared to GL and GBL. **(D)** Immunohistochemical staining against REELIN. Magnified Cajal-Retzius cells are indicated by arrows. Reduced numbers were seen in GB mice. In GBL mice Cajal-Retzius cells were located in deeper brain regions.

## Discussion

In this study we investigated the interplay of the developmental factor LIN28A and the WNT pathway key player CTNNB1, which are both described as oncogenic features in embryonic tumours with multi-layered rosettes (ETMRs) (Korshunov *et al*., 2012; Neumann *et al*., 2017; Lambo *et al*., 2019, 2020). An established mouse model for ETMR is initiated by co-activation of the WNT- and SHH- pathways but lacks the hallmark feature of LIN28A overexpression (Neumann *et al*., 2017). Therefore, we aimed to develop an ETMR mouse model displaying LIN28A overexpression. We show here that the co-activation of LIN28A and CTNNB1 in neural precursor cells during embryonic development - defined as the GBL model - was not sufficient to drive tumour formation. Instead, we observed histomorphological and molecular features resembling the human disorder Cobblestone lissencephaly type II (Devisme *et al*., 2012) with porous pial border and neuronal overmigration starting from E14.5. Using spatial proteomics, we show a shift in cortical layer identities with a change of the extracellular matrix (ECM) composition revealing new functions of LIN28A during brain development.

The RNA binding proteins LIN28A is an oncogenic factor being overexpressed in various cancer types occurring in the brain, ovary, breast, intestine or colon (Chen *et al*., 2011; King *et al*., 2011; Permuth-Wey *et al*., 2011; Lambo *et al*., 2019, 2020). The embryonal brain tumour ETMR expressing LIN28A also displays additional features, such as WNT and SHH pathway activation. Previous studies suggested that LIN28A expression acts upstream of these pathways within the developing tumour (Neumann *et al*., 2017; Lambo *et al*., 2019, 2020). However, a sole LIN28A overexpression in neural precursor cells does not lead to brain tumour formation (Middelkamp *et al*., 2021), suggesting that other factors are required to initiate tumour growth. CTNNB1 is an effector of the WNT signalling pathway which has been reported to be mutated in various cancers, including ETMR (Morin *et al*., 1997; Yokota *et al*., 2002; Neumann *et al*., 2017; Lambo *et al*., 2019, 2020). In our study, we showed that LIN28A alone and in combination with stabilised CTNNB1 is not sufficient to drive tumour formation during brain development. In other cancer types, such as intestinal cancer, LIN28A and its paralog LIN28B drive tumour formation, and cooperate with activated *Ctnnb1* (Tu *et al*., 2015), suggesting cell context dependent oncogenic functions. As a limiting factor, GBL mice did not survive postnatally, hence brain tumour formation at later developmental time points cannot be ruled out.

Previous studies investigating LIN28A overexpression demonstrated increased transient proliferation during early embryonic development (Yang *et al*., 2015; Middelkamp *et al*., 2021). Similarly, CTNNB1 has been described to increase the neural precursor pool during brain development by enhancing cell cycle re-entering and resulting in an enlarged cortical surface area and ventricle enlargement (Chenn and Walsh, 2002; Pöschl *et al*., 2013). At later embryonic time points, we detected a significant decrease of neural precursor cell proliferation in the GB model resulting in a thinned cerebral cortex, as described previously (Pöschl *et al*., 2013). Interestingly, the co-activation of both CTNNB1 and LIN28A (GBL) neutralised these effects and GBL mice showed no significant difference in proliferation compared to the CTRL. Respectively, GBL mice displayed a thicker cortex compared to GB mice but a histologically disturbed ventricular zone and cortical lamination. This might indicate a direct interaction between both factors. Of note, *LIN28A* was described to be regulated by the WNT pathway in breast cancer as a direct target of CTNNB1 (Cai *et al*., 2013). Additionally, CTNNB1 itself might be regulated at the mRNA level by LIN28A (Hafner *et al*., 2013) and reform its downstream function. Proliferative cells need sufficient supply ensured by biogenesis of required components. LIN28A promotes genes involved in ribosome biogenesis (Parisi *et al*., 2021) and dysregulation of ribosomal proteins is described in different cancer types supporting the proliferative behaviour (Grzmil and Hemmings, 2012; Elhamamsy *et al*., 2022). We showed that both LIN28A overexpressing GL and GBL models show dysregulated proteins belonging to the ribosomal family (Fig. 5), which have been described as potential LIN28A interaction partners (Hafner *et al*., 2013; Parisi *et al*., 2021). Nevertheless, the proteins associated with these terms were different in the GL and GBL model indicating context-dependent functional dynamics.

In addition to changes in progenitor cell proliferation, CTNNB1 activation was strongly related to migration disturbances and disordered lamination in the GB and GBL models. The disturbed migration was also represented in the proteomic signature. The GB model showed protein abundance changes in components involved in actin capping and filament assembly related to impaired cell motility (Fig. 5, Appendix Fig. S6) (Edwards *et al*., 2014). Similar to the *reeler* phenotype induced by knockout of the *Reelin* gene (Tissir and Goffinet, 2003), the GB model showed abnormal cortical lamination featured by reduced REELIN expression (Fig. 6B-D). Patients harbouring a *REELIN* mutation causing reduced Reelin protein abundance develop the migration disorder lissencephaly with cerebellar hypoplasia (LCH) (Hong *et al*., 2000). Generally, during development of the cerebral cortex, REELIN is mainly secreted by Cajal-Retzius (CR) cells in the marginal zone (MZ) regulating radial migration of neurons along the radial glia (RG) cells and fostering proper cortical lamination (Hirotsune *et al*., 1995; D’Arcangelo and Curran, 1998; Causeret *et al*., 2021). In contrast to the GB model, co-activation of CTNNB1 and LIN28A in GBL mice resulted in a severe migration disorder but increased REELIN expression was observed. Additionally, its receptor ITGB1 (Pinkstaff *et al*., 1999; Dulabon *et al*., 2000; Graus-Porta *et al*., 2001; Halder, Sapkota and Milner, 2022) was spatially dysregulated in the GBL model at E18.5. REELIN expressing CR cells responsible for proper radial migration were mis-localised in deeper cortical regions in the GBL model. The CR cells themselves are led along the meninges by secreted factors to the MZ (Borrell and Marín, 2006). At E14.5 we observed CR cells localised at the cortical surface indicating that CR cells reached their anticipated location, but disruption of the pial border and neural overmigration displace CR cells to deeper cortical regions. The neuronal migration disorder is known as the human disease Cobblestone (type II) lissencephaly (Devisme *et al*., 2012). Mutation in glycosyltransferases like FKRP and POMT1 (Brockington *et al*., 2001; Bouchet *et al*., 2007) lead to hypo-glycosylation of the DYSTROGLYCAN α-subunit (α-DYS) and thereby fail to interact with the extracellular matrix (ECM) resulting in a disturbed pial border (Barresi and Campbell, 2006). These glycosyltransferases were not detected in the proteome data, but we show that both CTNNB1 related models, GB and GBL have reduced α-DYSTROGLYCAN levels (Fig. 6B, C), indicating that this factor alone is not sufficient to disturb the pial border leading to neuronal overmigration.

To find additional components which explain the observed phenotype. We ablated layers from the skin surface into the cerebral cortex during embryonic development (E14.5 and E18.5) and aimed for comprehensive analysis of the spatially resolved proteome since LIN28A - as a RNA-binding protein - has the potential to alter the resulting proteome composition. This was also shown in LIN28A expressing ETMRs, where the proteome signatures significantly differ from their transcriptome profiles (Dottermusch *et al*., 2024). Layer signatures in the GBL model, revealed a lack of “positive regulation of cell-substrate adhesion” at E14.5 (Appendix Fig. S6). Later at E18.5, a shift in layer identity was observed. Superficial layers in the GBL model resembled deeper CTRL layers associated with the cerebral cortex indicating overmigration based on the proteomic signature. Additionally, high spatial deviation of the proteins related to the extracellular matrix (ECM) and their receptors was observed in the GBL model compared to the other models. The ECM protein LAMININ (LAMB1/LAMA1) (Hamill *et al*., 2009) was decreased corresponding to the disturbed pial border. The LAMININ receptor RPSA (Digiacomo and Meruelo, 2016) appeared as another interesting candidate which was dysregulated in the GBL model. It was shown that a knockdown of *Rpsa* leads to malformation of radial glia cells (RG) affecting the radial migration process (Blazejewski *et al*., 2022). This led to the hypothesis of improper integration of RGs and CRs due to accumulation of imbalances at the receptor and ligand level. In future studies it should be investigated how the altered molecular features fail the maintenance of the pial border and result in a severe form of overmigration in the GBL model (Appendix Fig. S8).

In summary, we show that in the GBL model LIN28A counterparts some effects of CTNNB1 preserving REELIN expression and stabilising the proliferation rate despite histological disturbances in the ventricular zone and cortical layering. Moreover, LIN28A affects translation of ribosomal components and components of the ECM revealing potential novel roles. Finally, the GBL model shows similarities with human lissencephaly type 2 and may be suitable for further analyses revealing the molecular foundations of this severe migration disorder.

## Methods

### Mouse models

The animals were kept at 12 h/12 h light/dark cycle with accessible water and food supply. Both male and female mice were examined. All experiments using animals were approved by the local animal care committee (Behörde für Lebensmittelsicherheit und Veterinärwesen in Hamburg) and handling was conducted in accordance with local governmental and institutional animal care regulations. The following mouse strains with C57BL/6 background were used: *hGFAP-cre* (Zhuo *et al*., 2001), *|S|- Lin28A(3x)-IRES-eGFP* (Papaioannou *et al*., 2013; Neumann *et al*., 2017; Middelkamp *et al*., 2021) and *Ctnnb1Δex3* (Harada *et al*., 1999; Pöschl *et al*., 2013). Breeding of the three respective mouse strains resulted in four different mouse models. Littermates containing floxed alleles but no *hGFAP-cre* were referred to as controls (CTRL). *hGFAP-cre::|S|-Lin28A(3x)-IRES-eGFP* (GL), *hGFAP- cre::Ctnnb1Δex3^FL/+^* (GB) and *hGFAP-cre::|S|-Lin28A(3x)-IRES-eGFP::Ctnnb1Δ ex3^FL/+^* (GBL) models were analysed during embryonic development at embryonic (E) days E14.5 and E18.5 (Appendix Fig. S1).

### Histology and immunohistochemistry

Formalin (4%) fixed and paraffin embedded tissue sections (2 µm) were used for Hematoxylin-Eosin (H&E) staining and for immunohistochemistry (IHC). The Ventana Benchmark XT machine (Ventana, Tuscon, USA) performed IHC with the following antibodies: BrdU (abcam, ab6326, 1:50), KI67 (abcam, ab15580, 1:100), MAP2C (Sigma-Aldrich, M4403, 1:3000), NeuN (millipore, MAB377, 1:100), pHH3 (Invitrogen Antibodies, MA5-15220, 1:100), REELIN (abcam, ab78540, 1:500), SOX2 (abcam, ab97959, 1:200), SSTR2 (abcam, ab134152, 1:1000), and TBR2 (abcam, ab23345, 1:250). Flash frozen tissue sections (8 µm) were used for IHC with LAMA1 (abcam, ab11575, 1:200). Digitalisation was performed with the Hamamatsu Photonics K.K. and NDP.view (version 2.8.24), and further image processing with Photoshop Elements 15 and Fiji (ImageJ 2.1.0, (Schneider, Rasband, and Eliceiri 2012)).

### BrdU injection and migration distance quantification

Pregnant mice at the embryonic (E) day E14.5 were injected intraperitoneally with 50 mg/kg BrdU (5-bromo-2-deoxyuridine), and the offspring sacrificed at embryonic day E16.5. Immunohistochemistry of frontal brain sections stained for BrdU and a semi-automated script for Fiji (ImageJ 2.1.0, (Schneider, Rasband, and Eliceiri 2012)) was used for the migration distance quantification of BrdU-positive (BrdU^+^) cells within the cerebral cortex. For the GBL model, we chose a region with clear ventricular border and thinned cortex for comparable analysis. For quantification, the scale was firstly calibrated for the distance measurement based on the image scale. Then the ventricular margin and BrdU^+^ cells were marked manually within the region of interest (ROI). Each distance was quantified automatically from the ventricular margin to the BrdU^+^ cell. To determine the migration distances across the mouse models, we defined ten bins each covering 50 µm and resulting in a range from 0 to 500 µm. The BrdU^+^ cells were assigned into the respective bin and a two-way ANOVA with Tukey’s multiple comparison test was performed. The cortex thickness was measured three times within each ROI for each replicate. The means of the cortex thickness and migration distances were compared using one-way ANOVA with Tukey’s multiple comparisons test. For all statistical test in GraphPad Prism (version 9.5.1, GraphPad Software Inc., La Jolla, California, USA): n = 3; p < 0.05.

### Proliferation

Immunohistochemistry of frontal brain sections with pHH3 at E14.5 and Ki67 at E18.5 was used for proliferation quantification. Using Fiji (ImageJ 2.1.0, (Schneider, Rasband, and Eliceiri 2012)), the ventricular margin was marked, and its length was measured. Positively stained cells (cells^+^) were counted along the marked ventricular margin and from the margin 50 µm into the ventricular zone. The margin length and cell count were used to calculate the proliferation rate = cells^+^/100 µm. One-way ANOVA with Tukey’s multiple comparison test (n ≥ 3, p < 0.05) was performed in GraphPad Prism (version 9.5.1, GraphPad Software Inc., La Jolla, California, USA).

### Western Blot

Lysis of fresh frozen tissues with 10% (w/v) homogenates in RIPA buffer (50 mM Tris–HCl pH 8, 150 mM NaCl, 1% NP-40, 0.5% Na-Deoxycholate, 0.1% SDS) freshly supplemented with 10x protease inhibitor and PhosStop (Roche, 04693159001) on ice. Total protein content was assessed by Bradford assay (BioRad, #5000205 and #5000206) and a total of 50 µg protein was loaded on the Any kD™ Mini-PROTEAN® TGX™ Precast Protein Gels (BioRad) or 4–12% Mini-PROTEAN® TGX™ Precast Protein Gels (BioRad). After electrophoresis separation, proteins were transferred to nitrocellulose membranes (BioRad, #1620213) by wet blotting. After washing, the membranes were blocked for 1 hour with 3% milk (ROTH, T145.2) in PBS for α-Dystrogylcan and β-Dystroglycan. 5% milk in TBST (TBS + 1% Tween20) was used as blocking buffer for the other antibodies. First antibodies: rabbit LIN28A (Cell Signaling Technology, 3978S, 1:250), mouse α-Tubulin (Developmental Studies Hybridoma Bank, 12G10, 1:2000), mouse α-Dystroglycan (Merck, 05-298, 1:100), mouse β-Dystroglycan (Sant Cruz Biotechnology, sc-165998, 1:400), mouse Reelin (abcam, ab78540, 1:500). The α-Dystroglycan and β-Dystroglycan antibodies were diluted with PBS, the other antibodies were diluted with the respective blocking buffer and incubated overnight at 4 °C. The blots were subsequently washed with PBS. Secondary antibodies: anti-mouse-IgG (Promega, W4028, 1:2500) and anti-rabbit-IgG (Promega, W401B, 1:2500) were diluted as the first antibodies and incubated for 2 hours at room temperature. Afterwards, washed with destilled water and TBST (TBS + 1% Tween20) and then incubated with the chemiluminescent reagent (Thermo Fisher Scientific, 34577) for visualisation. Protein abundance quantification was performed in ImageJ (ImageJ 2.1.0, (Schneider, Rasband, and Eliceiri 2012)). The values were normalised to the respective α-Tubulin and one-way ANOVA with Tukey’s multiple comparison test (n = 3, p < 0.05) was performed in GraphPad Prism (version 9.5.1, GraphPad Software Inc., La Jolla, California, USA).

### Samples and experimental setting for spatial proteomics

Using the nanosecond infrared laser (NIRL) nano-volume sampling system (Hahn *et al*., 2021; Voß *et al*., 2022), consecutive layers of about 40 µm thickness and an area of 800 µm x 800 µm targeting the forebrain as region of interest (ROI) were ablated and collected from fresh frozen material of embryonic mouse heads from the skin surface to the cortex for mass spectrometry-based characterisation with spatial resolution (Navolić *et al*., 2023). As reported before we ablated 9 layers for E14.5 CTRL mice (n = 4, 44 samples) (Navolić *et al*., 2023). In this work, samples of different mouse models were added at E14.5: GL (n = 3, 27 samples), GB (n = 3, 27 samples) and GBL model (n = 3, 26 samples). For E18.5 mice 18 layers were ablated for CTRL (n = 4, 81 samples), GL (n = 3, 54 samples), GB (n = 3, 54 samples) and GBL (n = 3, 54 samples). The samples D9 (E14.5, GBL) and I10-I18 (E18.5, CTRL) were excluded due to a shift in the region of interest (ROI) during sampling. Sample M4 (E14.5, CTRL) was excluded due to high abundance of human keratins. The plume of the ablated layer was collected and transferred in a tube (Protein LoBind® Tubes, Eppendorf SE, 0030108116) using 10 µl of 0.01 % DDM (n-dodecyl β-D-maltoside). For tryptic digestion 20 ng Trypsin was added. Technical parameters and all mass spectrometry preparation steps are described in further detail by Navolić et al., 2023 (Navolić *et al*., 2023).

### Mass spectrometry

The Tribrid Mass spectrometer (Orbitrap Fusion, Thermo Fisher Scientific, Waltham, MA, USA) was used for liquid chromatography-tandem mass spectrometer (LC-MS/MS) measurements coupled to a nano-UPLC (Dionex Ultimate 3000 UPLC system, Thermo Fisher Scientific, Waltham, MA, USA) in a data dependent acquisition (DDA) mode. Data base search of proteome data was performed with Proteome Discoverer (version 2.4.1.15). More details concerning the mass spectrometry parameters are described in the published work by Navolić et al., 2023 (Navolić *et al*., 2023).

### Data analysis

Spatial proteome data analysis was performed in RStudio (version 4.2.3). Of note, mass-spectrometry based data is a technique to investigate relative protein abundances, for simplicity we address it further only as protein abundances.

In total 367 samples were used for further analysis after the quality check (raw data: Table 1 and 2). The data was log2-transformed and normalised by median subtraction over samples. Each time point (E14.5 and E18.5) was further analysed separately.

Batch effect reduction was performed using the BERT framework (version 0.99.16, (Schumann and Schlumbohm 2024), in preparation). The mouse models and layers were integrated into classes serving as references. For E14.5: CTRL L1-L4, CTRL L5-L9, GL L1-L4, GL L5-L9, GB L1-L4, GB L5-L9, GBL L1-L4, GBL L5-L9; for E18.5: CTRL L1-L9, CTRL L10-L18, GL L1-L9, GL L10-L18, GB L1-L9, GB L10-L18, GBL L1-L9, GBL L10-L18 and corrected for the batch effect “laser ablation” (LA.batch). Correction for “measurement” (m.batch) was performed without references (data: Table 3). Dimension reduction analysis of the data was performed using Uniform Manifold Approximation and Projection (UMAP) (mixomics package, version 0.2.9.0) with proteins of 100% valid values, and visualised with GraphPad Prism (version 9.5.1, GraphPad Software Inc., La Jolla, California, USA).

ConsensusClusterPlus (version 1.62.0) was used to define layer clusters based on CTRL samples including proteins with missing values resulting in two clusters for E14.5: C_1_ (layer 1-4) and C_2_ (layer 5-9) and three clusters for E18.5: C_3_ (layer 1-2), C_4_ (layer 3-9) and C_5_ (layer 10-18). Ordering was changed without affecting the cluster results (Fig. 3D).

Differentially abundant proteins across layers were analysed with the limma package (version 3.54.2). For each comparison the replicates of a specific layer (L_n_) were compared to all other layers (L_all-n_) (i.e.: L _1_ vs L _2 to 9_) using proteins with 70% valid values (p < 0.05). For the CTRL model the top 10 high abundant proteins were selected for each layer. If a protein appeared in the consecutive layer it was excluded for all later layers resulting in the top 10 unique high abundant proteins per layer (first come, first serve principle) (Table 4). The unique CTRL layer signatures were then compared across the models layers by relative abundances using a heatmap and correlation analysis (corrplot (version 0.92)). The top high abundant proteins were also defined for the GL, GB and GBL model with the same approach.

Gene ontology (GO) analysis for biological processes (BP) was performed in EnrichR (version 3.2; (Chen *et al*., 2013; Kuleshov *et al*., 2016; Xie *et al*., 2021)) for every layer in all models using the significantly high abundant proteins (ordered by log Foldchange, max. 50 proteins). For every search the top 5 terms were selected. All terms were then further condensed using the rrvgo package (version 1.10.0, (Sayols, 2023)) to reduce redundant GO:BP-terms into parent GO:BP-terms. Afterwards, for each parent GO:BP-term a mouse model and layer deconvolution was performed. The “Best Match” value determines which layer in which model represents the parent GO:BP-term the best (highest –log10 of the mean adjusted p-values for each parent GO:BP term across layer and model (Table 5)). The “DiffScore” evaluates the difference of the represented layers across the models for each parent GO:BP-term. Firstly, the sum of layers for each model was calculated (i.e.: parent GO:BP-term_x_ in CTRL was represented in layer 4, 5 and 7: ∑ = 4+5+7 = 16). If a parent GO:BP-term was not present in a model is was assigned with the value -45 (S_9_ = - ([1+9)/2] *9)) in E14.5 or -171 (S_18_ = - ([1+18)/2] *18)) in E18.5. Then the range was calculated across mouse models (highest sum – smallest sum). A lower “DiffScore” indicates less change of layer representation across models, a higher value indicates a greater change or missing representation.

For each layer cluster (C_1_ - C_5_) an analysis of covariance (ANCOVA) was performed comparing proteins with 70% valid values in CTRL vs. another mouse model (GL, GB or GBL). Proteins with significant rate of change (slope) or abundance (intercept) were selected (p < 0.05 and R^2^ > 0.1) and categorised into specific proteins for the individual mouse model. These specific proteins were analysed with EnrichR (version 3.2; (Chen *et al*., 2013; Kuleshov *et al*., 2016; Xie *et al*., 2021)) for the top 5 GO:BP-terms for biological processes (GO:BP). The associated proteins were further categorised as potential interaction partners with LIN28A (line type): direct protein-protein interaction with LIN28A (Parisi *et al*., 2021) as mRNA targets of LIN28A (Hafner *et al*., 2013). Additionally, we used a literature miner pipeline (modified version (Steffen *et al*., 2020)) to evaluate the background knowledge of these proteins associated with the keyword “cortical development”. The proteins were assigned into three categories: 1 = at least one review article; 2 = at least one research article; 3 = no literature found (Steffen *et al*., 2020).

## Supporting information

Supplementary_Table_01_Sample and Protein annotation

Supplementary_Table_02_rawData_after quality check

Supplementary_Table_03_Data_norm_and_corrected

Supplementary_Table_04_top 10 unique proteins_CTRL

Supplementary_Table_05_parent GO_BP-terms

## Data availability

Full LC–MS/MS data are available via ProteomeXchange with identifier PXD053649.

Appendix: Fig. S1-S8 (pdf)

Supplementary Figures: Fig.S1-Fig.S8

Table 1: Supplementary_Table_01_Sample and Protein annotation (xlsx)

Table 2: Supplementary_Table_02_rawData_after quality check (xlsx)

Table 3: Supplementary_Table_03_Data_norm_and_corrected (xlsx)

Table 4: Supplementary_Table_04_top 10 unique proteins_CTRL (xlsx)

Table 5: Supplementary_Table_05_parent GO_BP-terms (xlsx)

## Acknowledgements

We would like to thank Tasja Lempertz, the Institute of Neuropathology and Kristin Hartmann from the Mouse Pathology facility for excellent technical assistance and service. We thank Hannah Voß for support with proteome analyses. We thank Ursula Müller, Jasmin Seydler, Alexandra Gröss, Anke Dorendorf, Beate Miksche and the other members of the animal facility team for their great support and service.

We used images and templates from http://smart.servier.com, licensed under a Creative Common Attribution 3.0 Generic License.

J.E.N was funded by the Deutsche Forschungsgemeinschaft (DFG, Emmy Noether programme, DFG project number 416054672). H.S. was supported by grants from the Deutsche Forschungsgemeinschaft (DFG) (INST 337/15-1, INST 337/16-1, INST 152/837-1, INST 152/947-1 FUGG and SCHL 406/21- 1).

## Author contributions

The manuscript was written by J.N. and J.E.N.. J.E.N. supervised and designed the study. S.H. and J.N. performed and interpreted western blots. M.Mi. conducted the BrdU experiments. M.D. developed the script for migration distance quantification performed by J.N.. J.N., M.Mo. and J.H. performed laser ablation experiments. M.Mo., C.K., H.S. performed proteome database search. J.N. and J.E.N. analysed and interpreted the data. J.N., Y.S. and A.G. performed data quality check and batch effect reduction.

A.G. modified literature miner tool (modified version (Steffen *et al*., 2020)). J.N., M.Mi., L.R., S.G. and P.S. generated and collected tissue samples. All authors reviewed the manuscript for intellectual content and approved the final version.

