## Supplementary Figures for "Co-activation of LIN28A and CTNNB1 disturbs cortical neuronal migration and pia mater integrity"

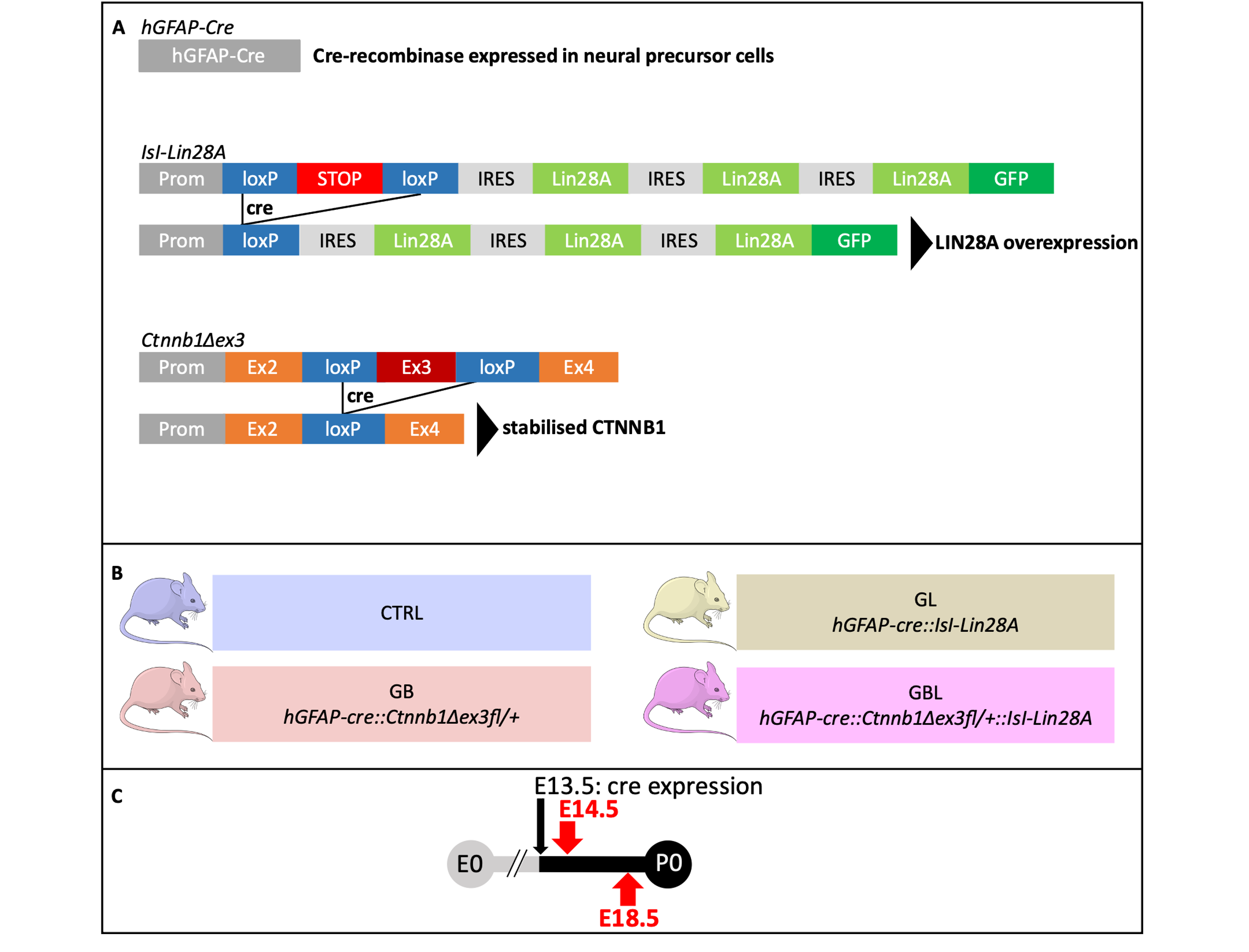
**Supplementary figure legends**

**Fig. S1: Scheme of transgenes and resulting mouse models.**

(A) Cre-Recombinase expressed in hGFAP-positive neural precursor cells leads to removal of a functional STOP sequence in the IsI-Lin28A construct. This results in LIN28A overexpression. Cre recombination of the Ctnnb1Δex3 construct removes exon3 resulting in stabilised CTNNB1. (B) Breeding of the three respective transgenic mouse strains resulted in four different mouse models: CTRL = control condition containing one or both floxed constructs but no Cre ( = no recombination event), GL = overexpression of LIN28A in hGFAP+ cells, GB = stabilised CTNNB1 in hGFAP+ cells, and GBL = overexpression of LIN28A and stabilised CTNNB1 in hGFAP+ cells. (C) hGFAP-dependent Cre recombination initiates at embryonic (E) day E13.5 (Zhuo et al. 2001). The time points E14.5 and E18.5 were investigated during development.

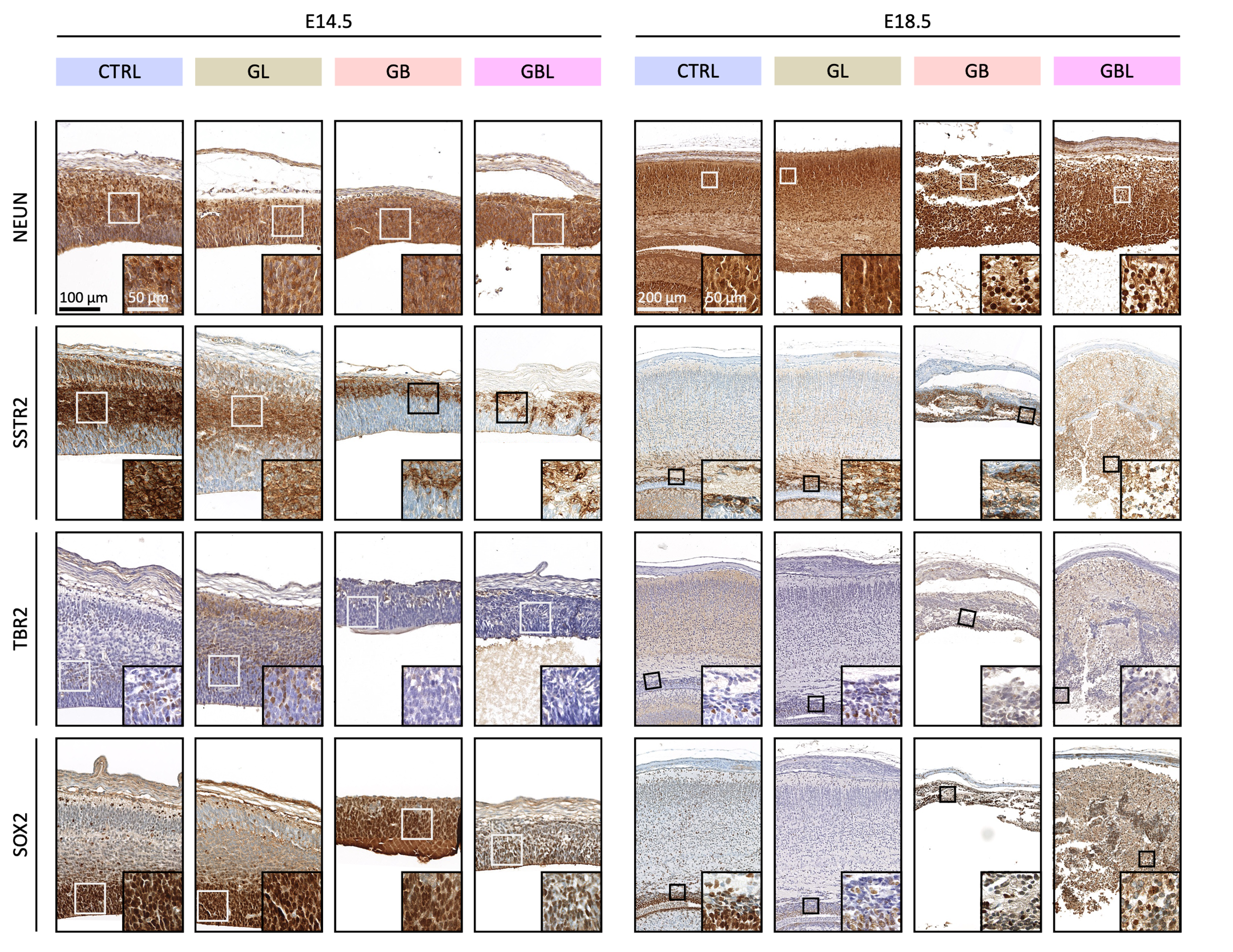

**Fig. S2: Cerebral cortex marker describing cortex layering.**

Immunohistochemical staining of cerebral cortices against NEUN, SSTR2, TBR2 and SOX2 at E14.5 and E18.5 in CTRL mice and the GL, GB and GBL mouse models. Scale indicates 100 µm. Magnified area is displayed in the bottom right corner of the respective image (scale in insets indicates 50 µm).

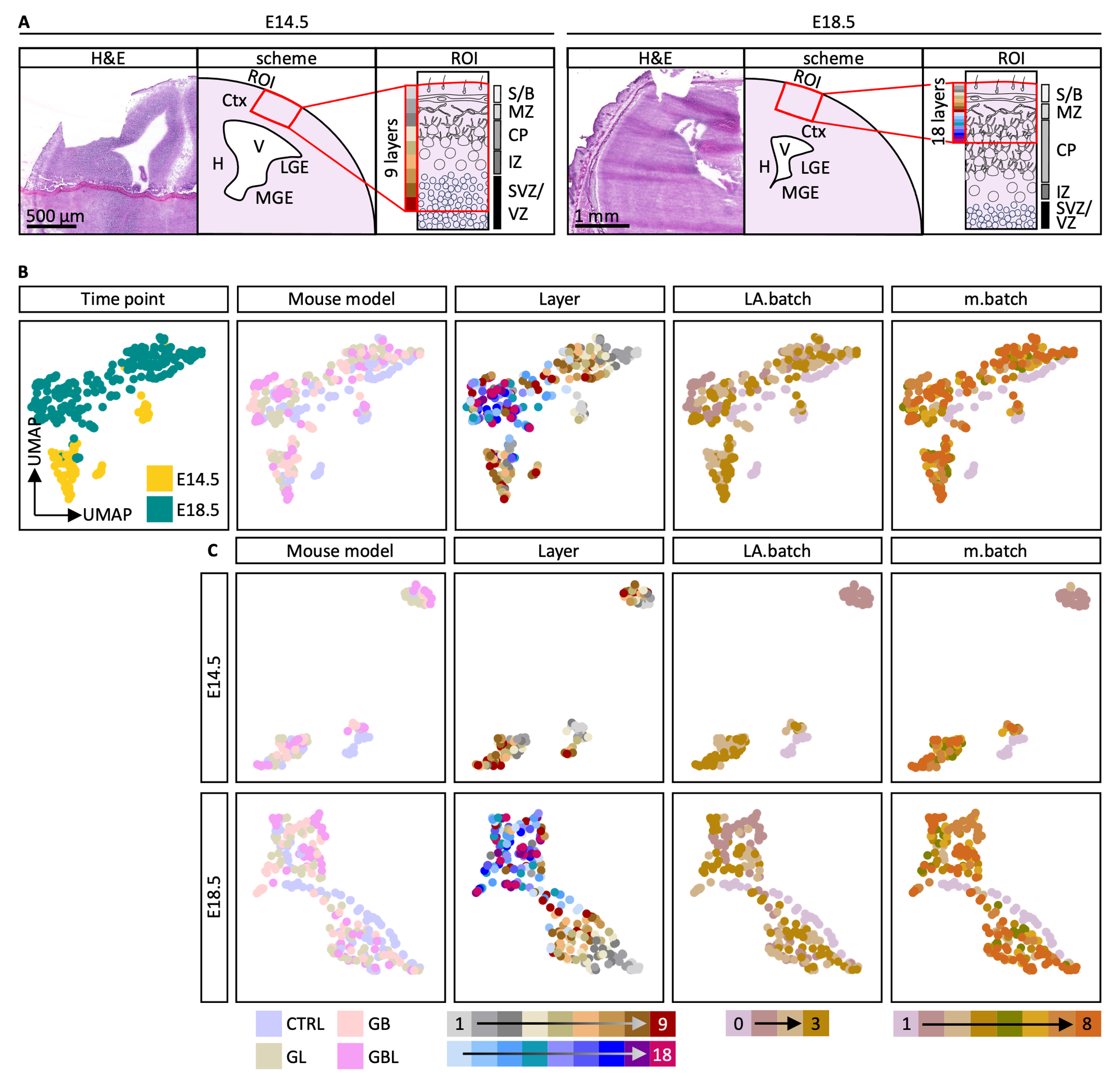

**Fig. S3: Sample overview for spatial proteome analysis.**

**(A)** Representative images of frontal H&E-stained sections at E14.5 and E18.5 after tissue ablation using NIRL. The scheme shows the region of interest (ROI) for ablation targeting the cerebral cortex (Ctx). Within the ROI, nine consecutive layers (thickness of each layer ~40 µm) were ablated from the skin into the cerebral cortex at E14.5. At E18.5 in total 18 consecutive layers were ablated. H = hippocampus, V = ventricle, MGE = medial ganglionic eminence, LGE = lateral ganglionic eminence, S/B = skin/bone, MZ = marginal zone, CP = cortical plate, IZ = intermediate zone, SVZ = subventricular zone, VZ = ventricular zone. **(B)** Uniform Manifold Approximation and Projection (UMAP) of all samples (n=367) before batch effect reduction based on proteins with 100% valid values. **(C)** UMAP displaying E14.5 and E18.5 data separately before batch effect reduction based on proteins with 100% valid values.

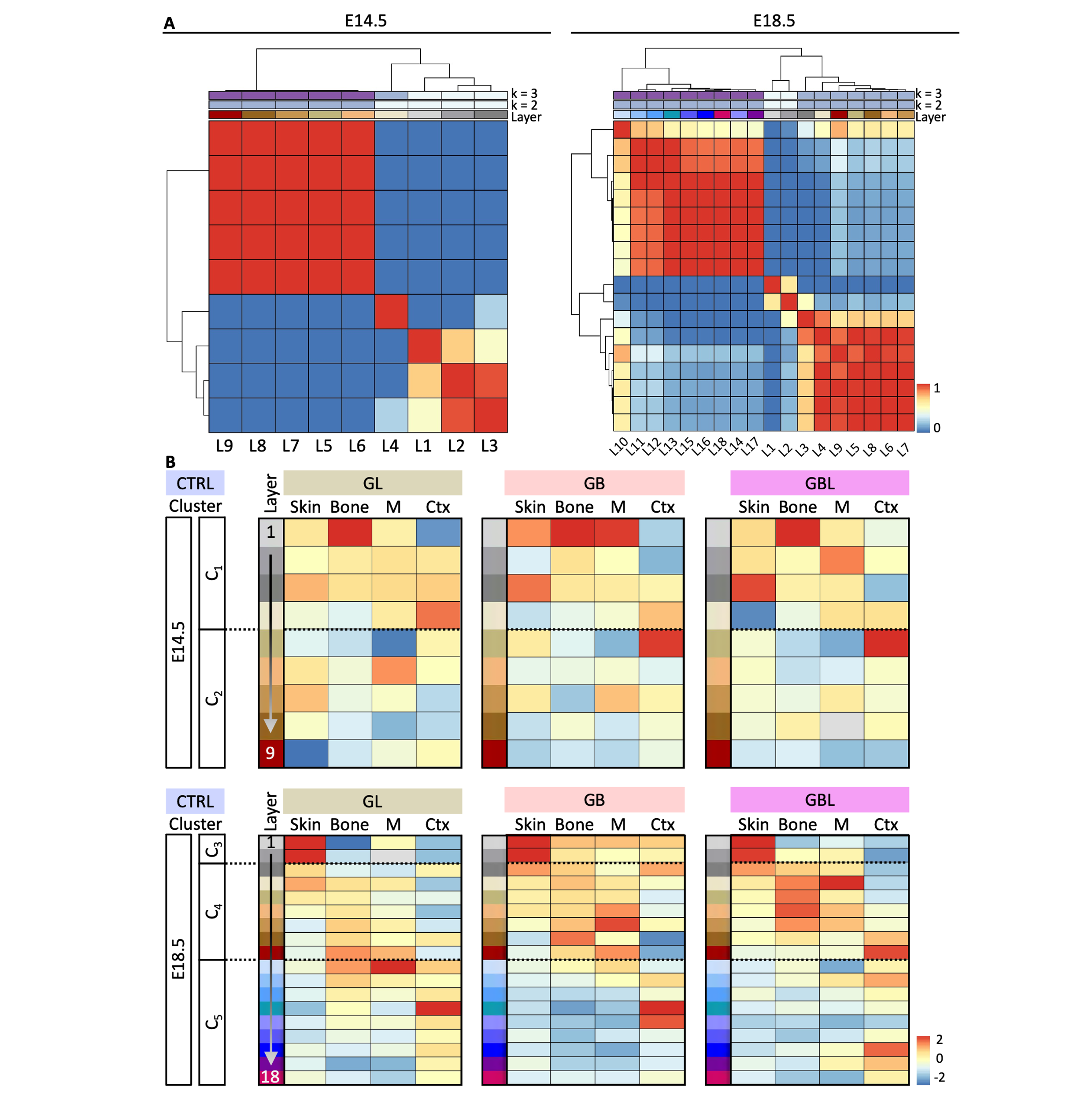

**Fig. S4: Layer clustering and profiling.**

**(A)** Heatmap of consensus clustering analysis of global proteome patterns in CTRL layers at time points E14.5 and E18.5. k = number of clusters, L = layer. **(B)** Mean column scaled abundance of marker proteins for Skin (FLG, KRT14, LORICIRIN), Bone (COL1A1, COL1A2, SERPINF1), M = Meninges (CDH11, CRABP2, TAGLN) and Ctx = cerebral cortex (TBR1, MAP2, BCL11B) in the respective mouse models GL, GB or GBL. C = cluster. C1-C5 represent clusters defined in **(A)** and are indicated with dashed lines.

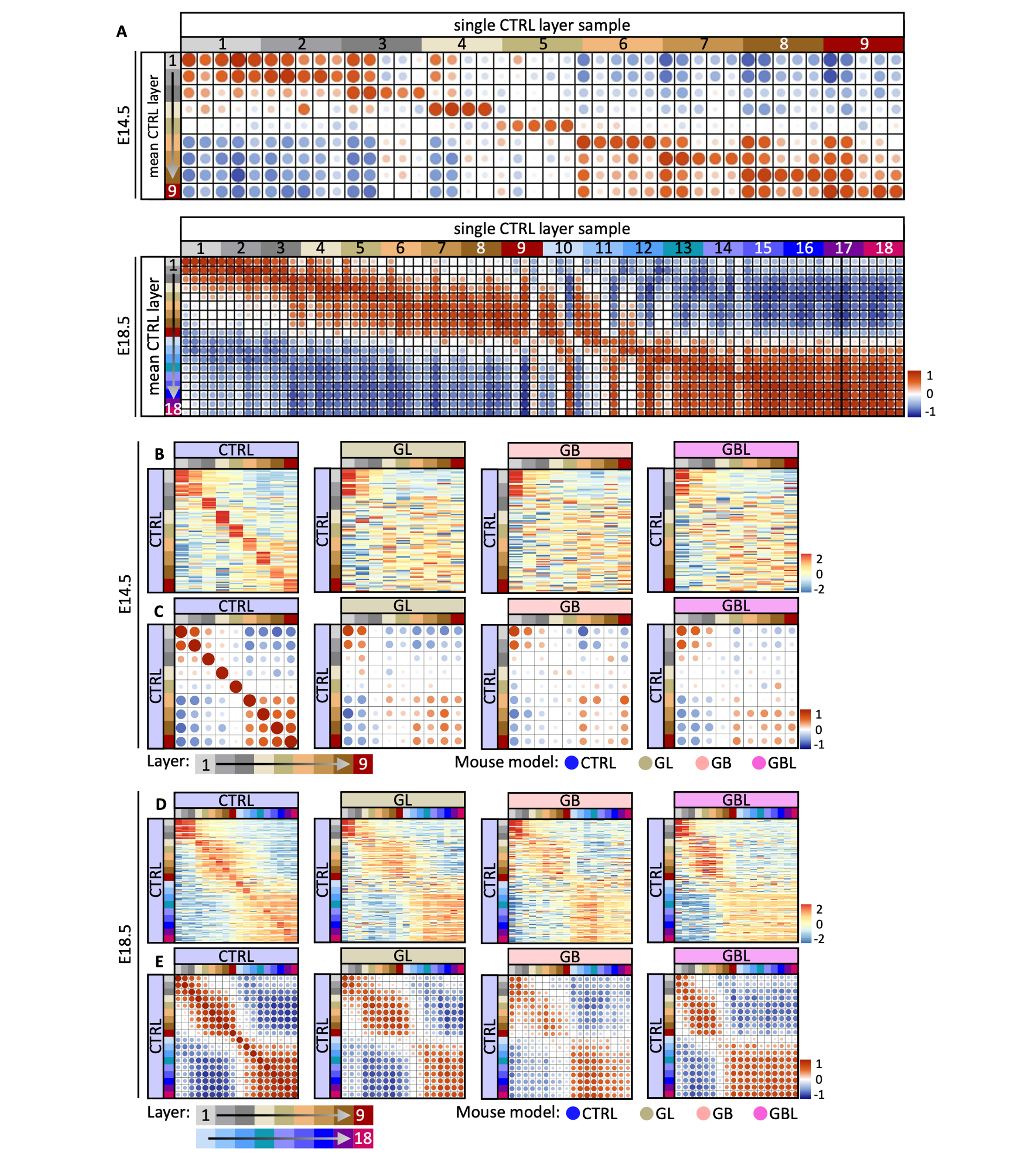

**Fig. S5: Layer signatures across mouse models compared to CTRL.**

**(A)** Correlation analysis based on the mean top 10 high abundant proteins of each CTRL layer displaying variation within the biological CTRL replicates. Protein abundances of each CTRL sample (x-axis) are correlated to the mean protein abundance of CTRL layers (y-axis). **(B, D)** Heatmap representing the top 10 high abundant proteins in each CTRL layer (y-axis) with row scaled abundances shown for CTRL, GL, GB and GBL layers (x-axis) at time point E14.5 **(B)** and E18.5 **(D)**. **(C, E)** Correlation analysis based on the top 10 uniquely high abundant proteins of each CTRL layer shown in **(B,D)**. Mean protein abundances of each CTRL, GL, GB and GBL layer (x-axis) are correlated with values of CTRL layers (y-axis).

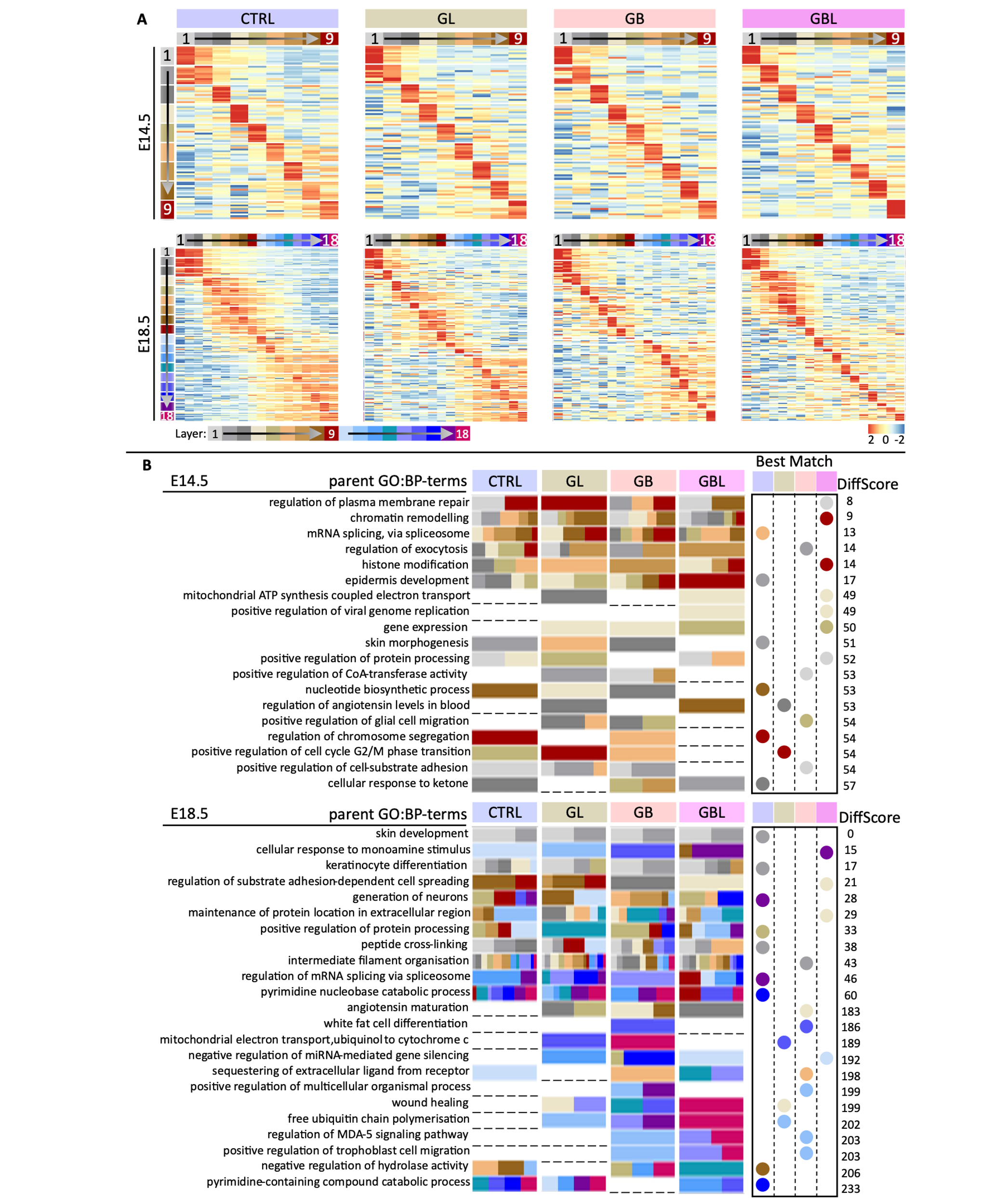

**Fig. S6: Unique layer signature for each mouse model and respective gene ontology (GO) analysis.**

**(A)** Heatmap representing the top 10 high abundant proteins of each layer for each mouse model (row scaled, Table 4). **(B)** Gene ontology GO analysis for biological processes (GO:BP) based on the top high abundant proteins of each layer for each mouse model. The top 5 terms of every search were further condensed into parent GO:BP-terms followed by mouse model and layer deconvolution. Blanks mean that the parent GO:BP-term was not represented in this mouse model. The Best Match shows which mouse model and layer (colour) represents the parent GO:BP-term the best based on adjusted p-values for each GO:BP-term (Table 5). The DiffScore indicates the (spatial) distance of representative layers for each parent GO:BP-term across mouse model. In E14.5 this score can range from 0 to 90 and at E18.5 0 to 342. The higher the value, the greater the difference of the represented layers across mouse models.

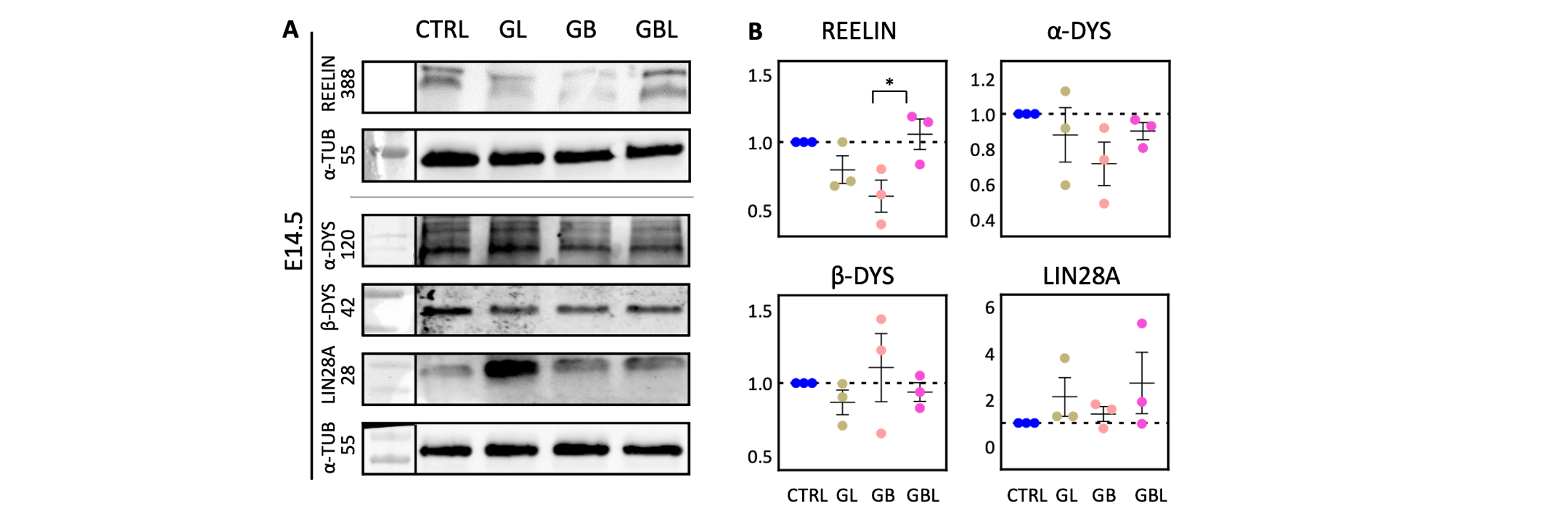

**Fig. S7: Extracellular matrix component**

**(A)** Western blot of cortical lysates stained against REELIN (388 kDa), α-DYS (α-DYSTROGLYCAN, 120 kDa), β-DYS (β-DYSTROGLYCAN, 42 kDa) and LIN28A (28 KkDa) at E18.5. α-TUB (α-TUBULIN, 55 kDa) was used as housekeeping protein for normalisation. **(C)** Respective quantification of western blot signals (B) based on normalised values (one-way ANOVA; n >= 3, * p < 0.05, ** p < 0.01, *** p < 0.001; mean and SEM are shown).

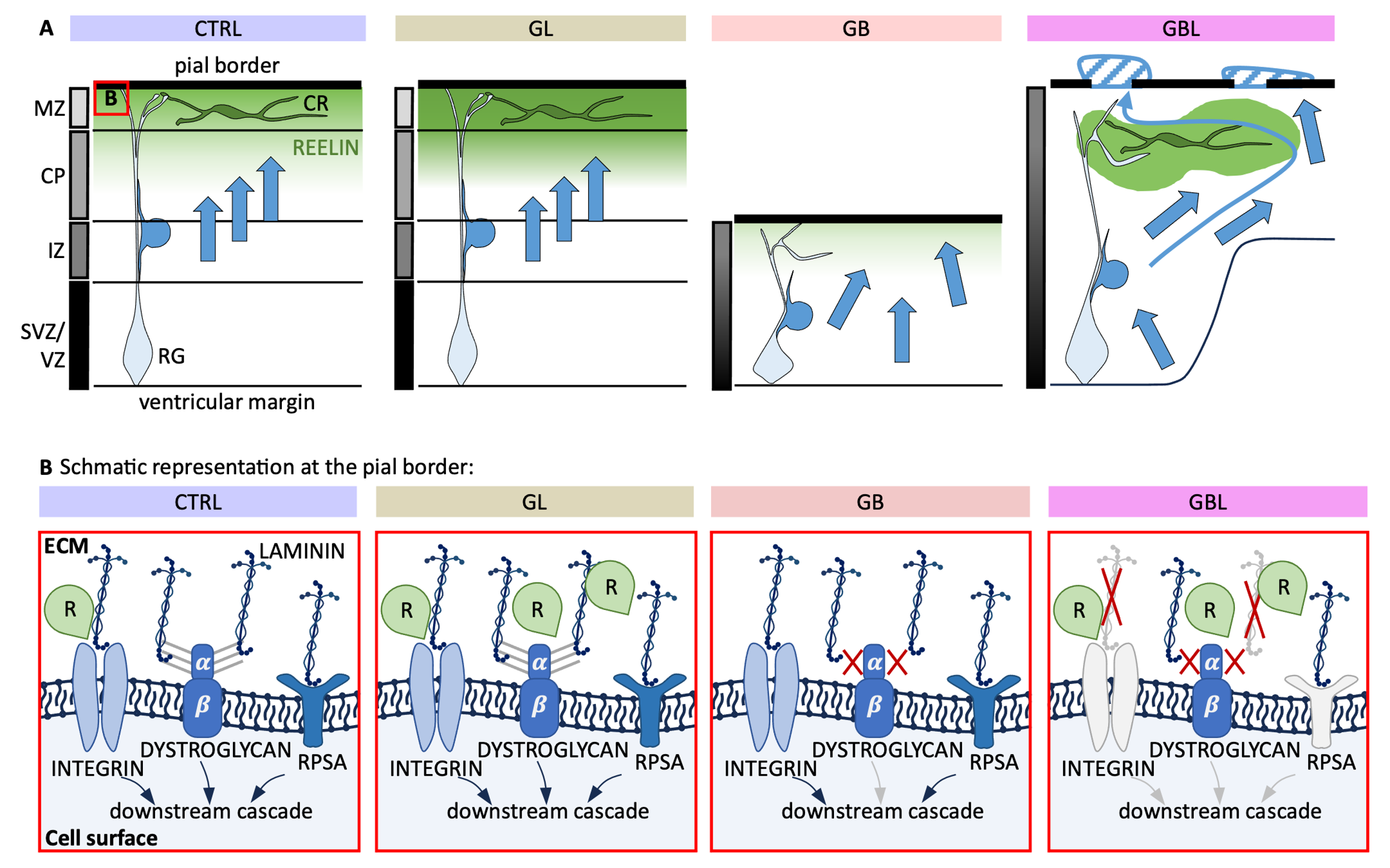

**Fig. S8: Summarising scheme of morphological and proteomic changes after LIN28A and CTNNB1 activation**

**(A)** CTRL condition showing a representative Cajal-Retzius (CR) cell in the marginal zone (MZ) and the radial glia (RG) cell as scaffold for radial migration of neuronal cells from the ventricular zone (VZ) through the intermediate zone (IZ) to their anticipated location in the cortical plate (CP) in an insight-out manner guided by a REELIN dependent (R) gradient. Red square indicates region at the pial border described in panel **(B).** In the GL model increased REELIN expression is seen with maintained cortical migration and lamination. The GB model showed reduced REELIN expression, malformation of RGs and disturbed migration with a failure to develop proper cortical lamination. The GBL model showed various cortical thickness, disturbed lamination and porous pial border with neuronal tissue ectopically located above the pial border. CR cells were located in deeper regions of the cerebral cortex, overexpressing REELIN. **(B)** LAMININ and REELIN are components of the extracellular matrix (ECM) and are recognised by their receptors at the cell surface. Binding to the receptors INTERGIN, DYSTROGYLCAN and RPSA induces downstream signalling relevant for migration and integration of neural cells in the cortical plate. REELIN levels were increased in the GL and GBL model. Hypo-glycosylation of α- DYSTROGYLCAN was seen in the GB and GBL model, and an accumulation of disturbed spatial expression of receptor components and LAMININ was detected in the GBL model.
